# Base Editing as a Therapeutic Strategy for Autosomal Dominant Tubulointerstitial Kidney Disease caused by Pathogenic *UMOD* Variants

**DOI:** 10.64898/2026.10.15.758759

**Authors:** Nora Diederich, Sarah Wendlinger, Shuhan Kong, Dimitra Papaioannou, Emily Haar, Amr Al-Shebel, Guy Yealland, Wenjun Jin, Carlos Cordero, Ria Schönauer, Roberta Di Costanzo, Marta Mariniello, Annika Fendler, Bernhard Ralla, Kai-Uwe Eckardt, Jan Halbritter, Olivier Devuyst, Verena Klämbt, Michael M. Kaminski

**Author notes:** Co-Corresponding authors Michael M Kaminski, Charitéplatz 1, 10117 Berlin, Verena Klämbt, Augustenburger Platz 1, 13353 Berlin.

## Abstract

Autosomal dominant tubulointerstitial kidney disease caused by pathogenic *UMOD* variants (ADTKD-*UMOD*) is an untreatable monogenic kidney disease in which misfolded uromodulin is retained in the endoplasmic reticulum and exerts a toxic gain-of-function effect. Here we systematically evaluate the full spectrum of base editing strategies targeting the *UMOD* locus. Base editors corrected a representative missense patient variant, p.Cys170Tyr, with A-to-G conversion efficiencies of up to 83.4%, restoring uromodulin trafficking to the plasma membrane in cell lines and in human 3D kidney tubuloids. Mutation-agnostic inactivation was achieved by four routes: premature stop codon introduction, splice-site disruption, allele-specific stop codon introduction and promoter attenuation. Maximum on-target efficiencies ranged from 44.1% to 71.9%, providing a strategy set applicable across the *UMOD* allelic spectrum. Endogenous *UMOD* expression is rarely recapitulated in human cell models, leaving strategies that act on the endogenous gene difficult to assess in patient cells. We therefore established CRISPR activation (CRISPRa) of the endogenous *UMOD* promoter and confirmed its ability to upregulate UMOD expression in human urine-derived cells. This approach further enabled quantification of endogenous *UMOD* transcript reduction following base editing in HEK293 cells. These results establish base editing as a versatile approach for ADTKD-*UMOD* and provide a primary human system for testing interventions targeting endogenous *UMOD* expression, from nucleic acid-based silencing to small molecules.

## Introduction

Autosomal Dominant Tubulointerstitial Kidney Disease (ADTKD) is a rare monogenic kidney disease characterized by tubulointerstitial fibrosis and progressive chronic kidney disease (CKD) in the absence of primary glomerular dysfunction^1^. The condition is genetically heterogeneous, resulting from pathogenic variants in *UMOD* (uromodulin), *MUC1* (mucin 1), *HNF1B* (hepatocyte nuclear factor 1B), *REN* (renin), or *SEC61A1* (translocon subunit α1)^1^. ADTKD caused by *UMOD* variants (ADTKD-*UMOD*) is the most frequent form of ADTKD, accounting for the third most common monogenic cause of CKD in adults, representing approximately 2–3% of familial CKD cases^2–4^. Uromodulin, encoded by *UMOD*, is the most abundant urinary protein, synthesized in the thick ascending limb (TAL) of the loop of Henle ^5^. It contains 48 evolutionarily conserved cysteine residues essential for correct folding and stability^5^, and serves multiple functions including regulation of sodium transport^5,6^, defense against urinary tract infections^5–7^, and prevention of kidney stone formation^8^.

Over 135 pathogenic *UMOD* variants have been described, predominantly heterozygous missense variants affecting cysteine residues or in-frame deletions. These variants induce a toxic gain-of-function effect likely mediated by protein misfolding and endoplasmic reticulum (ER) retention^9^, resulting in ER stress and unfolded protein response (UPR)^9–11^. This pathomechanism is supported by *Umod* knock-in mouse models recapitulating ER stress and the ADTKD-*UMOD* phenotype^12–18^.

In contrast, *Umod*-deficient mice exhibit no overt renal phenotype, except for increased susceptibility to urinary tract infections and loss-of-function variants in *UMOD* have not been reported to cause ADTKD in humans^19^. Beyond monogenic disease, common *UMOD* variants identified through GWAS are linked to increased risk of CKD^20^, hyperuricemia^21^, or hypertension^22^. These associations are partially attributed to enhanced uromodulin expression in the kidney, reflected by higher levels in urine^23^.

Clinically, patients with ADTKD-*UMOD* typically develop kidney failure (KF) around the fifth decade of life^1,19^. Once KF is reached, dialysis and kidney transplantation remain the only available options; however, dialysis is associated with increased morbidity and mortality, while transplantation is limited by organ shortage and the requirement for lifelong immunosuppression. Novel therapeutic strategies are therefore urgently needed.

Base editors have emerged as highly effective CRISPR/Cas-based tools capable of correcting single-nucleotide variants without DNA double-strand breaks, templates, or homology-directed repair^24^. Comprising a catalytically impaired nuclease fused to a deaminase, they currently enable A-to-G^24^, C-to-T^25^, A-to-Y^26^, and C-to-G^27^ edits, with preclinical efficacy demonstrated across diverse genetic diseases^28–32^. Recently, base editing has been used to correct *PKD1* variants in mouse models of autosomal dominant polycystic kidney disease (ADPKD), reducing both kidney and liver cyst burden^33,34^. Whether base editing is feasible for ADTKD-*UMOD* has not been explored.

Here, we test the full range of strategies base editing permits, from mutation-specific correction of individual *UMOD* variants to mutation-agnostic approaches applicable across the allelic spectrum. We assess rescue of the cellular disease phenotype in murine IMCD-3 (mIMCD) cells and in human 3D tubuloids derived from healthy nephrectomy tissue. As *UMOD* is not appreciably expressed in cultured human tubular cells, we propose using CRISPR activation to drive expression from the endogenous locus, enabling the mutant protein to be studied in the patient’s own genetic background.

## Material and Methods

### Generation and cultivation of HEK293T reporter and mIMCD-3 cell lines

HEK293T cell lines as well as HEK293/dCas9-VP64+MPH (CRISPR stable cell line HEK293/dCas9-VP64+MPH, SL302, GeneCopoeia Inc., Rockville, MD) cells were maintained in DMEM (Thermo Fisher Scientific, cat#41965062) supplemented with 10% fetal bovine serum (Sigma cat#F7524) and 1% P/S (gibco cat#15140122) at 37 °C with 5% CO_2_. mIMCD-3 cells stably expressing UMOD WT or C170Y were described previously^35^ and were maintained in Advanced DMEM/F-12 (Thermo Fisher Scientific cat#12634010) supplemented with 10% fetal bovine serum (Sigma cat#F7524), 1% L-Glutamine (GlutaMAX™ Supplement, Thermo Fisher Scientific cat#35050038) and 1% P/S at 37 °C with 5% CO_2_. HEK293T based cell lines were engineered to carry the target site (human *UMOD* C170Y, R185S or promoter variant rs4293393) by stable lentiviral integration or by using the sleeping beauty transposase system^36^. The variants, flanked by 150bp of surrounding genomic sequence, with specific viral primer sequences on the 5’ and 3’ ends, allowing for targeted sequencing, were synthesized (IDT gBlocks). Using Gibson assembly, the DNA fragments were either cloned into a lentiviral transfer plasmid with GFP tag (pWPXLd AG12258) or into pT4 SB plasmid (gift from Zsuzsanna Iszvak). Lists of primer sequences used are provided in **Supplementary Table S3 and S4**. HEK293T cells were transduced with the viruses at MOI 0.1 or electroporated with 4µg pT4 transposon plasmid and 1µg of the SB100x transposase RNA. After culturing the cells for 3-5 days, they were single-cell sorted for low GFP expression, and colonies of single clones were sequenced for the correct integration of targets.

### Generation and cultivation of human urine derived epithelial cells

Fresh spot urine samples from healthy donors were non-invasively collected at Charité Berlin after written informed consent (ethics vote under registration number EA4/066/21). Next, from urinary pellets, we generated urine derived renal epithelial cells (hURECs) according to a refined version^37^ of the Schutgens et al protocol^38^. hURECs were cultured in proximal tubular epithelial cell (PTEC) medium (DMEM-F12, Insulin 5 μg/mL, Thyroxin 5 μg/mL, Selenium 5 ng/mL, Hydrocortisone 36 ng/mL, EGF 10 ng/mL, Tri-iodothyronine 40 pg/mL, 10% FBS, 1% penicillin-streptomycin) as previously described^39^ for 2D culture. For tubuloid formation in 3D, hURECs were cultured for one passage in 2D in expansion medium (EM) ((ADMEM/F12 supplemented with 1% penicillin/streptomycin, HEPES, GlutaMAX, N-acetylcysteine (1 mM, Sigma) and 1.5% B27 supplement (Gibco)), supplemented with 10% Rspo1-conditioned medium, EGF (50 ng/ml, Peprotech), FGF-10 (100 ng ml/1, Preprotech), Rho-kinase inhibitor Y-27632 (10 µM, Abmole), A8301 (5 µM, Tocris Bioscience) and primocine (0.1 mg ml/1, Invivogen))^38^ and subsequently seeded in growth-factor-reduced Matrigel (Corning cat#356231).

### Generation and cultivation of renal tubuloids from biopsies

Primary renal cells were isolated from renal tissue obtained from patients undergoing nephrectomies due to renal carcinoma diagnosis as previously described^38^. Tissue collection was approved by the ethics committee of the Charite—University Hospital (EA1/274/22) and informed consent was obtained from all patients. Only healthy tumor-distant tissue was used for tubuloid generation. Fragments were seeded in growth-factor-reduced Matrigel and cultured as described above^38^.

### HEK293T and mIMCD-3 cell chemical transfection

HEK293T cell lines were seeded at low passages with a density of 1.25×10^5^ cells per 92µl in antibiotic-free medium in a 96-well cell culture microplate (Falcon cat#CLS351172) and grown for 24h at 37 °C with 5% CO_2_. Transfection with single-guide (sgRNA) and base editor (BE) plasmids was performed using the TransITX2 Dynamic Delivery System (Mirus Bio MIR cat#6004) following the manufacturer’s protocol in a 3:1 BE:sgRNA ratio by weight. 40ng total plasmid was transfected per well. 72h after transfection, gDNA isolation was performed. mIMCD-3 UMOD-GFP C170Y cells were seeded at low passages with a density of 3.125×10^5^ cells per 2500µl antibiotic-free medium in a 6-well cell culture plate (Corning cat#CLS3506) and grown for 24h at 37 °C with 5% CO_2_. Transfection with sgRNA and BFP-tagged BE plasmids was performed using the TransITX2 Dynamic Delivery System following the manufacturer’s protocol in a 3:1 BE:sgRNA ratio by weight. 1000ng total plasmid was transfected per well. 72h after transfection, cells were bulk-sorted for BFP expression using FACS and subsequently cultured for 72h at 37 °C with 5% CO_2_ in P/S-containing medium. Cells were then split for gDNA isolation and confocal imaging.

### Genomic DNA isolation

For HEK293T and mIMCD-3 cell line genomic DNA (gDNA) isolation, cells were washed once with PBS (DPBS, Gibco cat#14190250). mIMCD-3 cells were trypsinized (Trypsin-EDTA 0.25%, phenol red Gibco cat#25200072) and spun down at 250g for 4 min. HEK293T cell or mIMCD-3 cell pellets were resuspended in 70-100µl lysis buffer (100 mM Tris-HCl, pH 8.0, 200 mM NaCl, 5 mM EDTA, and 0.05% SDS), freshly supplemented with DTT and Proteinase K (25mM DTT, and 84 Units/ml Proteinase K NEB cat# P8107S). Samples were incubated for 16h at 55°C on a heatblock at 300rpm. In-house SPRI beads were prepared from Sera-Mag SpeedBeads Carboxylate-Modified Magnetic Particles, hydrophobic (Cytiva cat# 65152105050250) in DNA binding buffer (20% PEG-8000, 1.5 M NaCl, 10 mM Tris-HCl pH 8.0, 1 mM EDTA, 0.05% Tween-20), sterile-filtered (0.22 µm). gDNA was isolated and purified using these at a 1x ratio, with three washing steps using 80% ethanol and eluted in EB buffer. For quantification of gDNA, the Quant-iT dsDNA HS assay (Invitrogen cat#Q33267) was used.

### Targeted Amplicon Sequencing

Next-generation targeted amplicon sequencing (TAS) was performed to determine the efficiency of genome modification at the target sites using a two-step PCR-based library construction method adapted from the Illumina Nextera XT DNA library preparation. The target loci were amplified from 20-100ng of gDNA using the Q5 Hot Start high Fidelity 2x MM (NEB cat#M0494L) and PCR-1 primers. Primers with overhangs allowing for the subsequent indexing PCR for the engineered HEK293T cell lines were designed with PrimerBlast by NCBI for Stop introduction, splice site disruption and off-targets. Synthetic primers for use in HEK293T reporter cell lines were designed according to respective specific viral primer binding regions in HEK293T reporter cells. The PCR products were purified with the abovementioned SPRI beads at a 0.7x ratio and quantified using the Quant-iT dsDNA HS assay. Around 20-100 ng of purified PCR-1 products was used as template for the indexing PCR (PCR-2) to add barcodes and Illumina adapter sequences using Q5 and primers. PCR conditions for PCR-1 and PCR-2 are provided in **Supplementary Table S5**. PCR products were again purified using magnetic beads at 0.7x ratio, quantified and pooled at equimolar amounts. Pooled libraries were checked with the D1000 ScreenTape system (Agilent cat#5067-5582), spiked with 20-40% PhiX (Illumina cat#FC-110-3001), depending on the library complexity, and subsequently denatured. The final library was loaded on a MiniSeq sequencer at 1.5pM and sequenced as paired-end run using a Mini Seq Mid Output Kit (300 cycles) (Illumina cat# FC-420-1004). On-target genome editing activities were determined from sequencing data using CRISPResso2^40^ in pooled mode with custom input parameters for BEs: *min_reads_to_use_region: 100, quantification_window_size: 10, quantification_window_center:* −*10 base_editor_output– min_frequency_alleles_around_cut_to_plot 0*.*05*.

### mIMCD-3 cells immunofluorescence and image analysis

mIMCD-3 UMOD-GFP cells were grown in ibidi 8-well µ-slides (µ-Slide 8 Well, Ibidi cat#80826) for live imaging. Before imaging, cells were washed twice with DPBS containing Calcium and Magnesium (Gibco cat#14040117) and medium was replaced with phenol-red free culture medium (DMEM without phenol-red, Gibco cat#31053028, 10% FBS, 1% P/S) containing 1:10.000 nuclear stain (Thermo Fisher Scientific, Hoechst cat#33342, trihydrochloride, trihydrate cat# H1399) and 1:200 MemGlow™ 590 (Cytoskeleton, Inc. cat#MG03). After incubation for 12 minutes, confocal images were acquired using a Leica MICA confocal microscope with a 63×/1.20 water-immersion objective. Hoechst, EGFP and Cy3.5 were excited using 405, 488 and 561 nm laser lines, respectively. Identical acquisition settings were used for all samples within an experiment. Quantitative image analysis was performed by randomly selecting 10 visual fields for imaging from each replicate. Whole images were quantitatively analyzed.

### Protein Deglycosylation and Western Blotting

Protein extracts were obtained by dissolving cell pellets in Pierce RIPA buffer (Thermo Fisher Scientific cat #89900), incubation on ice for 30 min and centrifugation. For deglycosylation, 10-15 μg total protein lysate was denatured in 10× Glycoprotein Denaturing Buffer (New England Biolabs) at 100°C for 10 min and then cooled on ice. Samples were incubated with either Endo H (New England Biolabs) in 10× GlycoBuffer 3 or PNGase F (New England Biolabs) in 10× GlycoBuffer 2 supplemented with NP-40, at 37°C for 1 h. For SDS-PAGE, samples were mixed with Bolt LDS Sample Buffer (4×) (Thermo Fisher Scientific) and NuPAGE Sample Reducing Agent (10×) (Thermo Fisher Scientific), incubated at 70°C for 10 min and resolved on NuPAGE 3%–8% Tris-acetate gels (Thermo Fisher Scientific). Proteins were blotted onto PVDF membranes (Thermo Fisher Scientific) using the iBlot™ 2 dry blotting system (Thermo Fisher Scientific). Afterwards, membranes were incubated with antibodies against UMOD (ab207170, 1:1000; Abcam) or beta-actin (MA5-15739, 1:1000, Thermo Fisher Scientific), and secondary antibodies anti-rabbit IgG, HRP-linked Antibody (1:2000-10000, Cell Signaling cat#7074) or anti-mouse IgG, HRP-linked Antibody (1:5000, Cell Signaling cat#7076), respectively. Proteins were detected using Pierce ECL Plus Western Blotting Substrate (Thermo Fisher Scientific) and the intensity of individual immunoblot bands was determined by optical densitometry using ImageJ software (NIH). Uncropped western blot images are provided in the **Supplementary Fig. S3–S7**.

### Base editor plasmids and construct generation

ABEmax and BE4max were gifts from David Liu (Addgene plasmids #112095 and #112093, respectively), ABE8.20m was a gift from Nicole Gaudelli (#136300), and ABE8e and SaKKH-ABE8e were gifts from David Liu (#138489 and #138502). PAM-relaxed ABEmax constructs containing SpG, SpRY or SpCas9-NG (#140002, #140003 and #140005), as well as ABE8e- and ABE8.20m-based constructs containing SpG, SpRY or SpCas9-NG (#185911, #185912, #185914, #185916, #185917 and #185919), were gifts from Benjamin Kleinstiver. SpCas9 TadCBEd-V106W (#193839) and SpCas9-CBE6b (#215820) were gifts from David Liu. ABE9 was a gift from Dali Li (#194208). ABE9-derived SpG and SpRY constructs carrying P2A-tagBFP were generated in-house from the ABE9 backbone. All base editor plasmids used in this study are listed in **Supplementary Table S2**. Base editor plasmids obtained from Addgene were adapted for use in mIMCD-3 cells and HEK293/dCas9-VP64+MPH cells by substituting the eGFP reporter gene with tagBFP using a three-piece Gibson assembly. Expression plasmids for human U6 promoter-driven sgRNAs were generated by annealing and ligating duplexed oligonucleotides corresponding to spacer sequences into BsmBI-digested pUC19-U6-BsmBI_cassette-SpCas9_ sgRNA (BPK1520; Addgene plasmid #65777).

### GFP-Umod Plasmid

Genomic DNA (gDNA) was isolated from mIMCD-3 UMOD-GFP cells (WT and C170Y). The *UMOD-GFP* construct was amplified by PCR and cloned into the Sleeping Beauty pT4 transposon vector via Gibson assembly, incorporating a bovine growth hormone (bGH) poly(A) signal at the 3’ end. To generate lentiviral vectors for transduction of nephrectomy tissue-derived tubuloids, the respective constructs were subcloned from the Sleeping Beauty transposon plasmid into a pWPXLd lentiviral backbone via Gibson assembly. All primers used for Gibson assembly are provided in **Supplementary Table S3 and S4**.

### IVT RNA production

Linear DNA templates for in vitro transcription were generated from plasmid DNA by PCR using Q5 High-Fidelity 2x Master Mix (New England Biolabs). The forward primer was used to restore the functional CleanCap-compatible T7 promoter; the reverse primer introduced a 120-nucleotide poly(A) tail downstream of the 3′ UTR. PCR products were subsequently purified and used as templates for T7-mediated in vitro transcription. Transcription was performed in a total volume of 40 µL containing 1 µg linear DNA template, 5 mM each of ATP, GTP, CTP, and N1-methylpseudouridine-5′-triphosphate for reduced cytotoxicity, 4 mM CleanCap® Reagent AG, and T7 RNA polymerase mix. Following incubation at 37°C for 2h, RNA was purified using the RNA Clean & Concentrator™ Kit (Zymo Research), quantified, and stored at −80°C. Transcript size and integrity were assessed by agarose gel electrophoresis using an ssRNA ladder (New England Biolabs).

### RNA electroporation of tubuloids

Tubuloids were dissociated into fragments as previously described^38^ and subsequently treated with 0,25% Trypsin-EDTA at 37°C for 10 minutes. Single cell solution was electroporated with 5 µg of in vitro-transcribed RNA using P3 Nucleofector™ Solution in a 100-µL Nucleocuvette™ and the 4D-Nucleofector™ System (Lonza) with program DT-130. Following electroporation, cells were incubated for 3min at room temperature, recovered in culture medium (EM) and re-embedded in growth-factor-reduced Matrigel (Corning cat#356231) for subsequent three-dimensional culture in EM.

### CRISPR activation using Synergistic Activation Mediator (SAM)

For functional screening experiments in HEK293 cells, we employed a CRISPR stable cell line HEK293/dCas9 - VP64+MPH (SL302, GeneCopoeia Inc., Rockville, MD), expressing the SAM components. 10 sgRNAs binding on both the sense- and the antisense-strand up to 300bp upstream of the transcription start site (TSS) of UMOD were designed employing the online-tool CRISPick^41,42^. with the following settings for sgRNA selection: Target Gene ID: ENSG00000169344, Target Domain: [TSS-300,TSS+0], TSS Position: 20352715, PAM Policy: SpyoCas9. All sgRNA sequences are provided in **Supplementary Table S1**. For screening experiments in HEK293 cells, cells were subjected to antibiotic selection for expression of the SAM complex with blasticidin and hygromycin according to the manufacturer’s instructions. sgRNAs were cloned separately into the sgNRA (MS2) cloning backbone (addgene plasmid #61424). For sgRNA screening experiments, cells were seeded in 96-well plates at a density of 1.25 × 10^5^ cells/mL in a final volume of 92 µL per well. 24h after seeding, sgRNA-encoding plasmids were transfected using the TransIT-X2® Dynamic Delivery System, with a total of 40ng DNA per well. Each sgRNA was tested individually. In addition, the three best-performing sgRNAs identified in the initial screening were co-transfected at a 1:1:1 mass ratio, also totaling 40ng. Cells were harvested 56h after transfection for RNA isolation and subsequent qPCR analysis, as detailed below. Based on these screening experiments, act_sgRNAs1, 2, and 9 were selected for all subsequent CRISPRa experiments and combined at an equal mass ratio (hereafter referred to as act_sgRNA mix). For functional evaluation of splice site disruption and stop codon integration in HEK293/dCas9-VP64+MPH cells, cells were seeded in a 6-well format at a density of 2.5 × 10^5^ cells/mL in a final volume of 2500µl per well. 24h after seeding, transfection with sgRNA and BE plasmids was performed using the TransITX2 Dynamic Delivery System in a 3:1 BE:sgRNA ratio by weight with 1250µg plasmid in total. For splice site disruption functional readout with qPCR, cells were then re-seeded at a density of 1.25 × 10^5^ cells/mL in a final volume of 92µl per well in a 96-well format. 24 hours after seeding, cells were transfected using 40µg act_sgRNA mix. Lysis for RNA isolation was performed 56h after transfection. For Stop codon introduction functional evaluation with WB, 72h after transfection, cells were re-seeded in a 6-well format at a density of 2.5 × 10^5^ cells/mL in a final volume of 2500µl per well. After 72h, cell pellets were harvested for western blotting.

### RNA electroporation of hURECs

For CRISPRa in hURECs, plasmid DNA was delivered by electroporation using P3 Nucleofector™ Solution, 100-µL Nucleocuvette™ vessels, and the 4D-Nucleofector™ System (Lonza) with program DT-130. A total of 5 × 10^5^ cells were electroporated per sample with 5µg plasmid DNA comprising dCas9-VP64 (Addgene plasmid #61422), MS2-p65-HSF1 (Addgene plasmid #61423), and the act_sgRNA mix at a 1:1:1 mass ratio. Within the act_sgRNA mix, the three sgRNAs were present at an equal 1:1:1 mass ratio. 56h after transfection, cell pellets were harvested for western blotting.

### qPCR

Gene expression was quantified directly from cell lysates using the Luna® Cell Ready Probe One-Step RT-qPCR Kit (New England Biolabs cat#E3031) according to the manufacturer’s instructions. Cells were lysed in 40 µL of lysis mix per sample, and RT-qPCR was performed using a CFX96 Real-Time PCR Detection System (Bio-Rad). Relative gene expression was calculated using the 2^−ΔΔCt method and normalized to ACTB expression. RT-qPCR conditions and primer and probe sequences are provided in **Supplementary Table S6 and S7**.

### Statistical analysis

Statistical analysis was performed using GraphPad Prism version 8.4.3. Data are presented as mean ± SD unless otherwise indicated. Statistical analyses were performed on independent biological experiments, with technical replicates averaged before statistical testing. Comparisons between two groups were performed using two-tailed unpaired Student’s t-tests or, for matched experiments, two-tailed paired Student’s t-tests. Comparisons between multiple groups were performed using one-way ANOVA followed by Dunnett’s or Tukey’s multiple-comparisons test, as appropriate. P values <0.05 were considered statistically significant. Statistical significance is indicated as *P*<0.05, *\*P*<0.01, *\*\*P*<0.001 and *\*\*\*P*<0.0001. Exact n values and the statistical tests used for individual experiments are indicated in the respective figure legends.

## Results

### Correcting the pathogenic UMOD variant p.C170Y by base editing

Pathogenic *UMOD* variants cluster in exon 3 and commonly affect conserved cysteine residues^19^. We first investigated base editing for correction of the well-characterized p.C170Y variant, which shares both features and impairs uromodulin maturation and trafficking (**Fig. 1**)^35^. To evaluate base editing *in vitro*, we generated a HEK293T reporter cell line via lentiviral integration of a sequence harboring the *UMOD* p.C170Y variant flanked by 150 bp of surrounding endogenous *UMOD* sequence. We then delivered various combinations of base editors and sgRNAs via transient transfection and assessed editing efficiency and precision through targeted amplicon sequencing (TAS) (**Fig. 1A**). For the p.C170Y variant, we identified four suitable single guide RNAs (sgRNAs) differing in their protospacer sequences and PAM requirements (**Fig. 1B**), which were experimentally tested using three ABEs (ABEmax^43^, ABE8.20m^44^, ABE8e^45^) in combination with four Cas9 variants (SpCas9, SpCas9-NG, SpG, SpRY^46,47^) for expanded PAM recognition, resulting in a total of 15 compatible combinations. Editing efficiencies for the intended A-to-G conversion ranged from 6.7% to 82.8%, with the ABE8e–SpG / C170Y_sgRNA2 combination achieving the highest efficiency (**Fig. 1C**). Importantly, cells were not pre-selected for transfection success, which likely led to an underestimation of true editing efficiency. We then assessed editing outcomes for potential bystander mutations **(Fig. 1D-E**) and identified a bystander edit at position A11 (in reference to C170Y_sgRNA2 A11>G), resulting in an Asp-to-Gly substitution at amino acid position 172, p.Asp172Gly (**Fig. 1D**).

**Figure 1.**
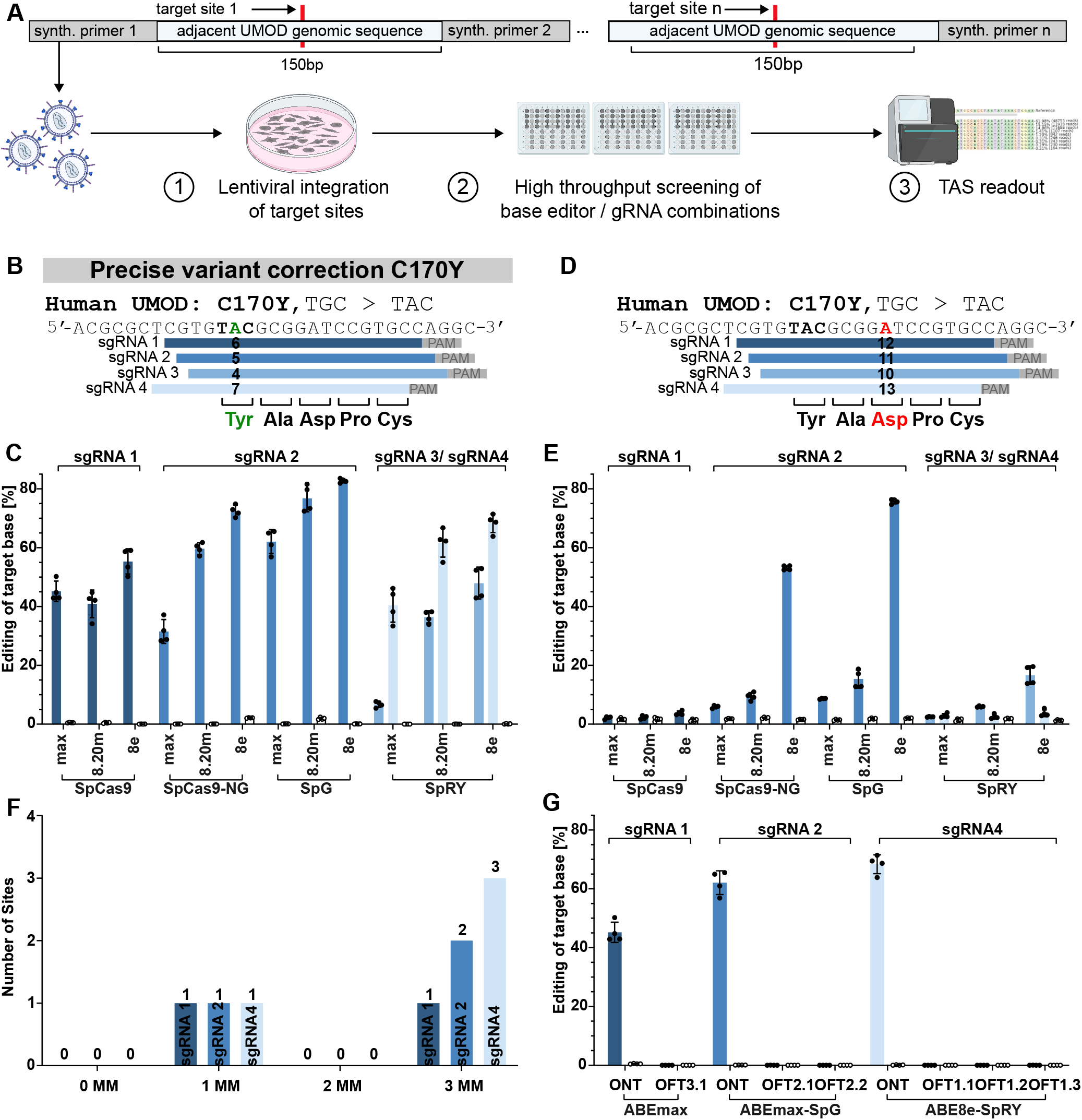
Base editing enables efficient and precise correction of the UMOD p.C170Y variant. (**A**) Experimental workflow: target sites with 150 bp of endogenous flanking sequence were introduced into HEK293T cells by lentiviral transduction, followed by transfection with base editor and sgRNA combinations and targeted amplicon sequencing (TAS). (**B**) sgRNAs targeting the UMOD p.C170Y variant (C170Y_sgRNA1–4, shown as sgRNA1–4), with sgRNAs (blue bars), PAM sequences (grey boxes), target base (green) and the tyrosine-encoding triplet (bold). (**C**) On-target editing efficiencies for the indicated sgRNA-base editor combinations, using ABEmax (max), ABE8.20m (8.20m) and ABE8e (8e) coupled to SpCas9, SpCas9-NG, SpG or SpRY. (**D**) Positions of bystander adenines for the UMOD p.C170Y variant within the same four sgRNAs, shown as in (B). The main bystander adenine is highlighted in red, with its position within each protospacer indicated. All nucleotide substitutions within the sgRNA target sequence, except for the intended edit, were considered bystander edits. (**E**) Bystander editing efficiencies for the combinations in (C). (**F**) Cas-OFFinder-predicted off-target sites for sgRNA1 with SpCas9-ABEmax, sgRNA2 with SpG-ABEmax and sgRNA4 with SpRY-ABE8e (up to three mismatches (MM), no bulges). y-axis, number of predicted sites; x-axis, number of mismatches. (**G**) On- and off-target editing efficiencies for the three combinations in (F), determined from the same samples. Predicted off-target sites were tested for C170Y_sgRNA1 with ABEmax (OFT3.1), C170Y_sgRNA2 with ABEmax-SpG (OFT2.1–2.2), and C170Y_sgRNA4 with ABE8e-SpRY (OFT1.1–1.3). (**C, E, G**) y-axis: percentage editing of the target base (C,G) or bystander bases (E) determined by TAS 72 h after transfection. CTL, base editor without sgRNA (white bars). Bars show mean values, dots individual technical replicates (n = 4), error bars standard deviation; experiments were independently repeated (n = 2). sgRNAs are shown in shades of blue throughout. sgRNAs tested are indicated above brackets at the top of the graph.

This bystander edit was most prominent with ABE8e-SpG and ABE8e-SpCas9-NG in combination with C170Y_sgRNA2 (**Fig. 1E**). Replacing ABE8e with the less active ABE8.20m or ABEmax reduced, but did not eliminate, bystander editing across all sgRNAs tested. Using C170Y_sgRNA4 shifted the bystander adenine to position 13, minimizing detectable bystander editing, but required the near-PAMless SpRY variant. Notably, combining NGG-dependent SpCas9 with C170Y_sgRNA1 enabled efficient on-target editing without detectable bystander editing, offering a high-specificity editing strategy. In this configuration, the target adenine was positioned at nucleotide 6, while the bystander adenine was shifted to position 12, largely outside the editing window. These results highlight the importance of optimizing sgRNA positioning, PAM compatibility, and base editor permissivity as well as editing windows to minimize bystander editing while maintaining efficient on-target editing.

As these combinations targeting the p.C170Y locus may differ in genome-wide specificity, we combined computational *in-silico* prescreening and experimental validation to identify potential off-target sites. Using the web-tool Cas-OFFinder^48^, we screened for predicted off-target sites of the three most promising combinations C170Y_sgRNA4 and ABE8e-SpRY, C170Y_sgRNA2 and ABEmax-SpG and C170Y_sgRNA1 with ABEmax-SpCas9 with up to 3 mismatches compared to the on-target site (**Fig. 1F**) as well as up to 1 RNA or DNA bulge (**Supplementary Fig. S1**). Sequencing of all identified sites revealed no increase in mutation rates in base editor treated versus untreated samples, demonstrating high on-target specificity with no detectable off-target editing under the conditions tested (**Fig. 1G**). Taken together these results demonstrate efficient and precise editing of the C170Y locus through systematic evaluation with the most favorable profile achieved through combination of C170Y_sgRNA1 and ABEmax–SpCas9, especially due to the more conservative editor choice without detectable bystander or off-target editing.

### Installing the protective rs4293393-G promoter variant

As pathogenic *UMOD* variants act through a toxic gain-of-function mechanism, lowering uromodulin expression may represent a mutation-agnostic strategy applicable to a broader spectrum of patients. We first targeted the *UMOD* promoter, leveraging the naturally occurring promoter variant rs4293393-G, carried by ~19% of Europeans. The G allele is associated with reduced *UMOD* transcription and a protective effect against CKD, in contrast to the more common rs4293393-A risk allele, which is linked to elevated uromodulin levels, increased CKD risk and hypertension^20,23^. We therefore designed a base editing strategy to convert the rs4293393-A allele to G, thereby mimicking the naturally protective haplotype (**Fig. 2A**).

**Figure 2.**
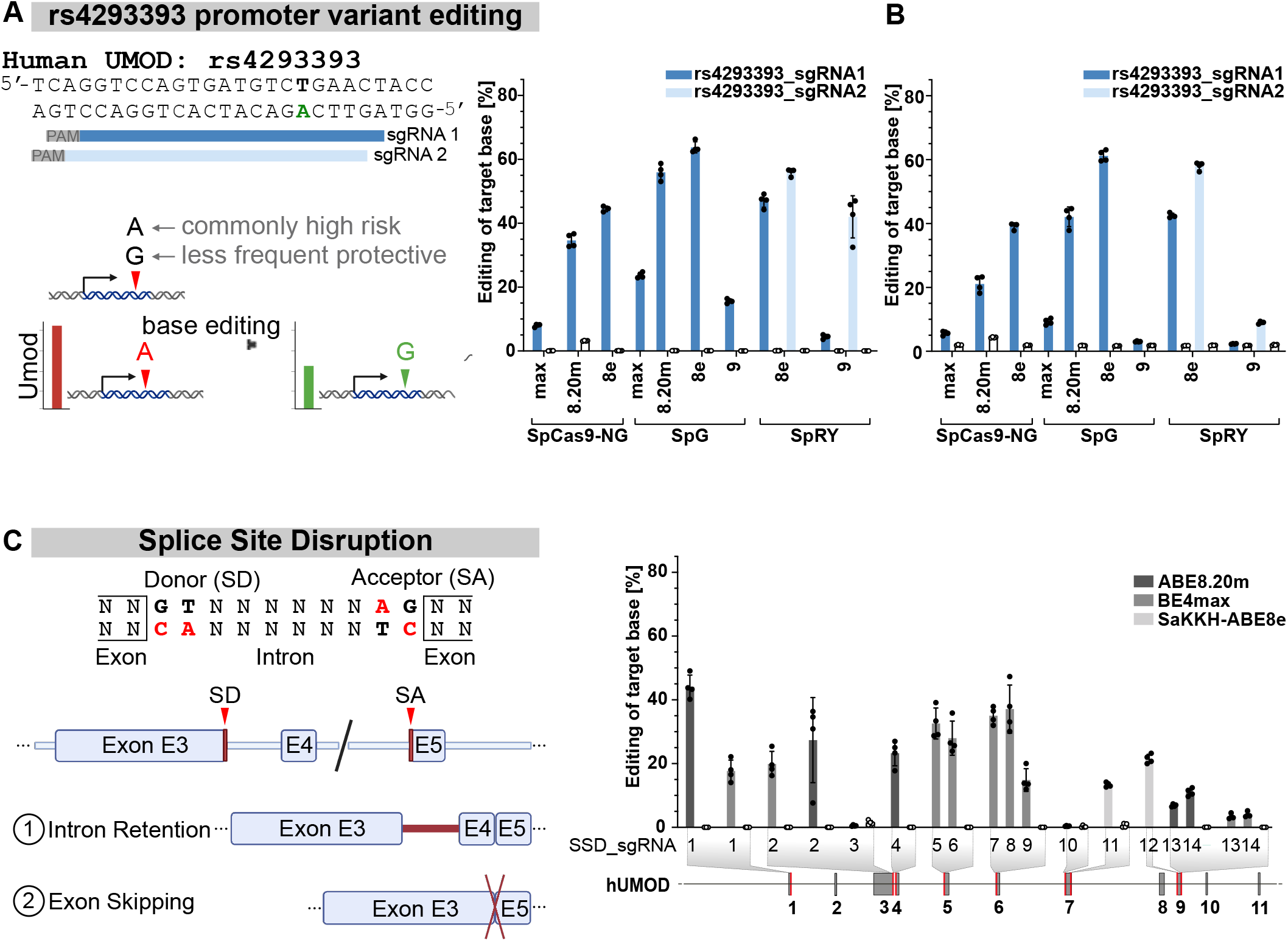
Mutation-agnostic base editing of the *UMOD* promoter and essential splice sites. (**A**) Left: schematic of editing the naturally occurring rs4293393-A risk allele in the *UMOD* promoter to install the protective rs4293393-G variant associated with reduced *UMOD* expression, as illustrated below. sgRNAs (blue bars), PAM sequences (grey boxes), target base (green) are indicated. Right: editing efficiencies at rs4293393 using the ABEmax, ABE8.20m, ABE8e and ABE9 deaminases in combination with SpCas9-NG, SpG and SpRY Cas9 variants, as indicated on the x-axis. Bars are shaded according to the suitable respective sgRNA tested, as indicated in the legend. (**B**) Bystander editing efficiencies for the sgRNA-base editor combinations shown in (A). All nucleotide substitutions within the sgRNA target sequence, except for the intended edit, were considered bystander edits. (**C**) Left: schematic of splice-site disruption (SSD) for functional gene inactivation by adenine or cytosine editing of conserved splice donor and acceptor sites, leading to intron retention (1) or exon skipping (2), as illustrated. Right: editing efficiencies at *UMOD* splice sites for SSD_sgRNA1–14, tested with SpCas9-based ABE8.20m and BE4max or with SaKKH-ABE8e, as indicated in the legend. sgRNA locations are indicated below the x-axis on the *UMOD* transcript. (**A, B, C**) y-axis: percentage editing of the target base (A,C) or bystander bases (B) determined by TAS 72 h after transfection. CTL, base editor without sgRNA (white bars). Bars show mean values, dots individual technical replicates (n = 4), error bars standard deviation; experiments were independently repeated (n = 2).

Two sgRNAs were designed (rs4293393_sgRNA1 and rs4293393_sgRNA2), with the target base located at position 6 and 5, respectively. rs4293393_sgRNA1, having an NG PAM, was tested with three ABEs (ABEmax, ABE8.20m and ABE8e) harboring the two Cas9 variants SpCas9-NG and SpG as well as ABE8e-SpRY. rs4293393_sgRNA2, having an NR PAM, was tested with ABE8e-SpRY only. This overall yielded eight PAM-compatible combinations. We achieved an A-to-G conversion efficiency of up to 64% at the rs4293393 locus in a HEK293T reporter cell line (**Fig. 2A**). Additional A-to-G conversions were observed at positions 8 and 10 for rs4293393_sgRNA1 and positions 7 and 9 for rs4293393_sgRNA2 (**Fig. 2B**).

Due to the observed unwanted bystander editing, both sgRNAs were subsequently also tested using ABE9^49^, for rs4293393_sgRNA1 carrying a SpG or SpRY Cas9 variant and for rs4293393_sgRNA2 carrying a SpRY Cas9 variant. Due to the positioning of the target adenine at protospacer positions 6 and 5, respectively, in both cases the target falls within the narrow editing window of ABE9. Notably, rs4293393_sgRNA2, with the target base at position 5, showed comparatively high on-target editing efficiency (42.0%) while bystander editing was significantly lowered at 9.0%, which is consistent with the highly restricted deamination window of ABE9. Among the tested combinations, ABE9-nSpRY with rs4293393_sgRNA2 thus provided the most favorable ratio between on-target editing and additional nucleotide conversions. These results demonstrate efficient installation of the protective rs4293393-G allele at the *UMOD* promoter, establishing a mutation-independent strategy that recreates a naturally occurring human haplotype rather than introducing a novel sequence change.

### Disrupting essential UMOD splice sites

Whereas promoter editing installs an allele associated with reduced *UMOD* transcription, we next asked whether base editing could inactivate the gene completely, an approach supported by the tolerance of *UMOD* loss of function in mice. We first targeted essential splice sites, selecting 14 sgRNAs pre-screened using SpliceR^50^ and testing them with SpCas9-based ABE8.20m and BE4max^43^, which require an NGG PAM, or with SaKKH-ABE8e^45^ (**Fig. 2C**), which recognizes an NNNRRT PAM and is smaller in size at around 3.9kb, potentially facilitating in vivo delivery by reducing vector payload requirements. Editing efficiencies ranged from 0.4% to 44.1% across all combinations. The highest on-target editing was achieved at the GT splice donor site of *UMOD* intron 1 with SSD_sgRNA1 and ABE8.20m (44.1%), followed by the AG splice acceptor site of intron 6, targeted with BE4max and SSD_sgRNA7 or SSD_sgRNA8 (35.0% and 37.1%, respectively) (**Fig. 2C**). Compared with precise variant correction, occurrence of bystander edits in gene inactivation may be less consequential, as these are unlikely to compromise the intended disruption. The effect of these splice-site disrupting edits on *UMOD* transcript levels was assessed in the cell systems described below. Cas-OFFinder screening results for off-targets are shown in **Supplementary Fig. 1B**. Together, these results show that essential *UMOD* splice sites can be edited efficiently with both adenine and cytosine base editors.

### Introducing premature stop codons into UMOD

As a second route to inactivation, we introduced premature termination codons directly. We screened 12 positions within endogenous *UMOD* in HEK293T cells at which cytosine base editors could convert a tryptophan or glutamine codon into a stop codon by C-to-T transition (in exons 2-8) (**Fig. 3A**). To introduce premature stop codons, sgRNAs were designed adjacent to NGG PAM sequences and evaluated using three cytosine base editors: BE4max, an established APOBEC1-based CBE; CBE6b^51^, an evolved TadA-derived base editor with enhanced C-to-T selectivity and reduced unwanted A-to-G editing, a potential cause of stop codon loss; and TadCBEd-V106W^52^, which has been reported to exhibit a narrower editing window and substantially reduced Cas-independent DNA and RNA off-target activity. On-target editing efficiencies ranged from 7.5% to 71.2% across all combinations, with the highest efficiency achieved using STOP_sgRNA4 and TadCBEd-V106W (**Fig. 3A**). Editing efficiencies showed similar target-site-dependent trends across all three editors. However, CBE6b generally performed similarly to BE4max, while TadCBEd-V106W achieved comparable or higher efficiencies than BE4max and CBE6b at most target sites, suggesting that its reported improvements in specificity do not necessarily compromise on-target activity. Cas-OFFinder screening results for off-targets are shown in **Supplementary Fig. 2**. Together, these results identify suitable target sites for efficient installation of premature stop codons in *UMOD* and highlight CBE6b and especially TadCBEd-V106W as promising alternatives to BE4max.

**Figure 3.**
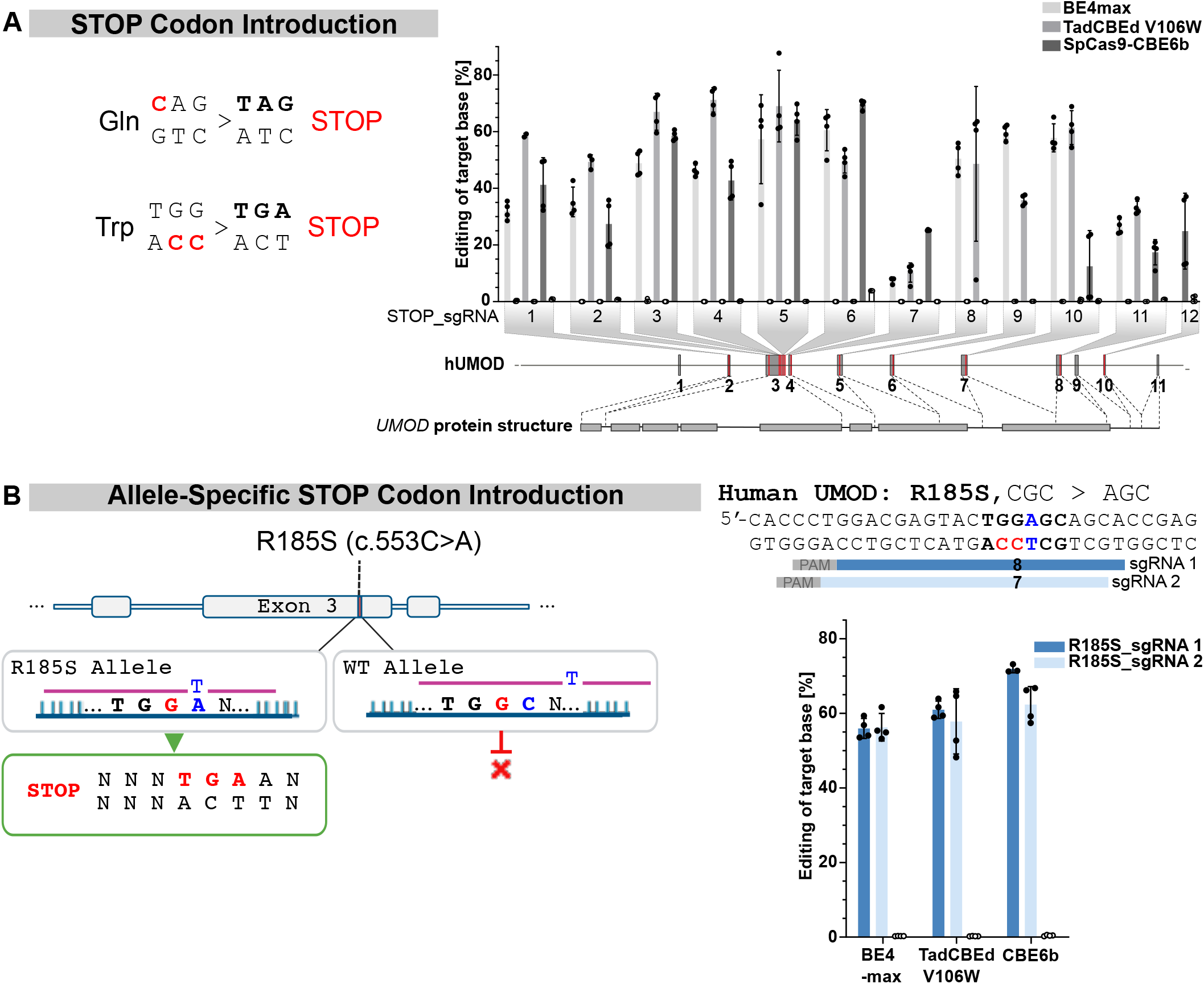
Introduction of premature and allele-specific stop codons into *UMOD*. (**A**) Left: Introduction of premature in-frame stop codons for gene silencing by cytosine base editing. Right: Editing efficiencies for STOP_sgRNA1–12 targeting tryptophan or glutamine codons within endogenous *UMOD*, tested with BE4max, TadCBEd-V106W and SpCas9-CBE6b as indicated in the legend. Guide positions along the *UMOD* transcript are indicated in red on the schematic below. (**B**) Left: Schematic of allele-specific stop codon introduction at the p.R185S pathogenic variant for selective silencing of the disease-conferring allele. The pathogenic variant lies within the protospacer, enabling preferential targeting of the mutant allele (green box, successful stop codon introduction) while sparing the WT allele (red cross, no editing). Right, top: schematic of editing the R185S pathogenic. sgRNAs (blue bars), PAM sequences (grey boxes), target bases (red) for stop codon introduction and allele-specificity conferring R185S variant (blue) are indicated. Right, bottom: On-target editing efficiencies for allele-specific stop codon introduction at p.R185S using BE4max, TadCBEd-V106W and SpCas9-CBE6b with R185S_sgRNA1 and R185S_sgRNA2, as indicated in the legend. (**A, B**) y-axis: percentage editing of the target base determined by TAS 72 h after transfection. CTL, base editor without sgRNA (white bars). Bars show mean values, dots individual technical replicates (n = 4), error bars standard deviation; experiments were independently repeated (n = 2).

### Introducing an allele-specific premature stop codon into the mutant UMOD allele

Whether preserving the wild-type allele is required for therapeutic benefit in ADTKD-*UMOD* remains an open question. In a mouse model, wild-type uromodulin mitigated aggregation of the mutant protein^35^, as proposed for other dominant proteinopathies, though this has not been shown in patients. We therefore asked whether the sequence context created by a pathogenic variant could itself be exploited for allele-specific stop codon introduction, using p.R185S, which is associated with earlier-onset kidney failure and more rapid progression than p.C170Y^35^. The p.R185S substitution (CGC→AGC) lies within the protospacer, so that sgRNAs matching the mutant sequence carry a mismatch against the wild-type allele, conferring inherent allele specificity (**Fig. 3B**).

We designed R185S_sgRNA1 and R185S_sgRNA2 targeting sequences adjacent to NGG PAMs, positioning the target cytosine at protospacer positions 8 and 7, respectively. C-to-T editing of the non-coding strand results in the corresponding G-to-A substitution on the coding strand, thereby generating a premature stop codon (TAG/TGA/TAA). Once again, both sgRNAs were evaluated with the SpCas9-based editors BE4max, TadCBEd-V106W, and CBE6b in a HEK293T reporter cell line (**Fig. 3B**). Editing efficiencies ranged from 56.0% to 71.9% across all combinations, with the highest achieved using R185S_sgRNA1 and CBE6b (**Fig. 3B**). Cas-OFFinder screening results for off-targets are shown in **Supplementary Fig. 1C**. These results show that pathogenic variants creating a discriminating protospacer sequence can be exploited for allele-specific inactivation of *UMOD*.

### Establishing cellular models of the UMOD trafficking defect

Having established efficient editing at the DNA level, we next sought to develop cellular models in which the functional consequences of base editing could be assessed. As standard cell lines do not express *UMOD*, we generated two complementary models expressing wild-type or mutant human uromodulin. We first used mIMCD-3 cells (murine inner medullary collecting duct), a well-established renal tubular epithelial line that has previously been used to study uromodulin trafficking and the ER retention of pathogenic variants stably expressing GFP-tagged wild-type (WT), p.C170Y or p.R185S hUMOD^35^ (**Fig. 4A**). Plasma membranes were labeled with MemGlow and GFP-membrane co-localization was assessed by confocal microscopy (**Fig. 4B**). WT hUMOD showed a predominant GFP signal at the plasma membrane, whereas both p.C170Y and p.R185S displayed a diffuse cytoplasmic signal (**Fig. 4B**), reflected in a significantly reduced Manders’ colocalization coefficient from 0.48 in WT to 0.20 and 0.29 for p.C170Y and p-R185S, respectively (**Fig. 4C**). To confirm that this reflects retention within the ER, we assessed the glycosylation state of GFP-tagged WT, p.C170Y and p.R185S uromodulin by digestion with endoglycosidase H (Endo H), which cleaves high-mannose glycans acquired in the ER but not the complex glycans acquired in the Golgi, and with PNGase F, which removes all N-linked glycans (**Fig. 4D**). Wild-type UMOD exhibited both Endo H-sensitive immature and Endo H-resistant mature forms, consistent with passage through the Golgi, whereas C170Y- and R185S-UMOD was predominantly Endo H-sensitive, with no detectable Endo H-resistant fraction, confirming ER retention (**Fig. 4D**).

**Figure 4.**
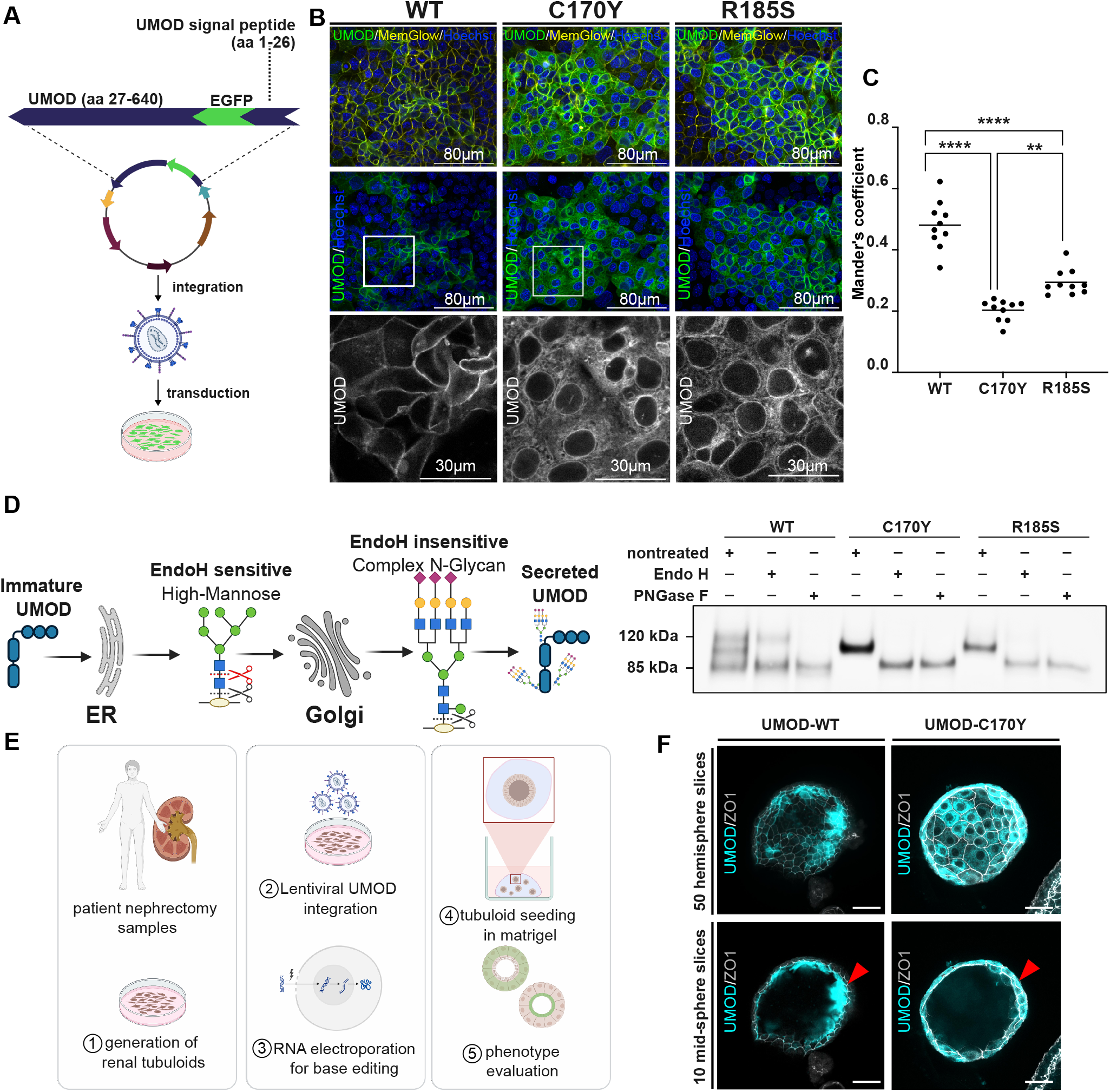
Cellular models of the *UMOD* trafficking defect. (**A**) Schematic of the lentiviral construct encoding GFP-tagged human uromodulin (hUMOD) and its stable integration into mIMCD3 cells. (**B**) Representative confocal images of mIMCD-3 cells stably expressing GFP-tagged WT, p.C170Y or p.R185S hUMOD (green), with plasma membranes labeled using MemGlow (yellow) and nuclei counterstained with Hoechst (blue). Scale bars, 80 µm (overview images) and 30 µm (magnified images). (**C**) Quantification of GFP-membrane colocalization by Manders’ colocalization coefficient for WT, p.C170Y and p.R185S hUMOD. Individual dots represent data from individual images. Horizontal bars indicate mean values from n = 10 images; ordinary one-way ANOVA with multiple comparisons,*\*\*P* < 0.0027, *\*\*\*\*P* < 0.0001 (**D**) Left: schematic of uromodulin N-glycan maturation along the secretory pathway. High-mannose glycans acquired in the ER are converted to complex glycans in the Golgi; Endo H (red) cleaves high-mannose but not complex glycans, whereas PNGase F (black) removes all N-linked glycans. Protein retained in the ER therefore remains Endo H-sensitive, while protein that has transited the Golgi becomes Endo H-resistant. Right: western blot of GFP-tagged WT, p.C170Y or p.R185S hUMOD in mIMCD-3 cells after mock treatment, Endo H digestion or PNGase F digestion. (**E**) Schematic of experimental workflow for base editing in nephrectomy-derived renal tubuloids. 1. tubuloids are established from nephrectomy samples, 2. lentiviral integration of *UMOD* constructs, 3. base editor mRNA and sgRNA are delivered by RNA electroporation, 4. edited cells were reseeded in matrigel for 3D tubuloid formation and 5. subsequent phenotypic analysis. (**F**) Representative immunofluorescence images of human 3D tubuloids expressing GFP-tagged WT or p.C170Y hUMOD (cyan) and stained for zonula occludens-1 (ZO-1, light grey). Scale bars, 50 µm. Red arrowheads indicate apical release of Umod into the tubular lumen for *UMOD*-WT tubuloids and intracellular retention of Umod for *UMOD*-C170Y tubuloids.

In tubular epithelium, uromodulin is GPI-anchored, sorted apically and released into the lumen^53^. To examine this in a human, tissue-derived context, we used tubuloids derived from adult human kidney, which span multiple nephron segments, remain genomically stable over many passages and form a polarized epithelium enclosing a lumen^38^.

As in other kidney organoid systems, endogenous *UMOD* expression was insufficient, such that we stably introduced GFP-tagged WT or p.C170Y h*UMOD* before 3D seeding (**Fig. 4E**). WT uromodulin localized apically and accumulated within the tubuloid lumen, whereas p.C170Y uromodulin was retained intracellularly (**Fig. 4F**), confirming that the trafficking defect seen in 2D is reproduced in a polarized human epithelium. These models therefore reproduce the cellular phenotype of ADTKD-*UMOD* in both a 2D system suited to quantitative screening and a polarized human 3D epithelium.

### Rescuing the p.C170Y trafficking defect by base editing in mIMCD-3 cells

Having demonstrated efficient and precise correction of p.C170Y at the DNA level, we next asked whether base editing rescues the associated cellular phenotype. p.C170Y mutant hUMOD mIMCD-3 cells were co-transfected with the sgRNA and a blue fluorescent protein (BFP) labeled ABEmax, enabling enrichment of successfully transfected BFP-positive cells isolated by FACS. Compared with cells transfected with the base editor alone, base-edited cells showed a pronounced redistribution of GFP signal from intracellular compartments to the plasma membrane, closely resembling the WT localization pattern (**Fig. 5A**). This was reflected in the Manders’ colocalization coefficient, which averaged over three biological replicates reached 0.44 in edited p.C170Y cells, comparable to 0.48 in WT cells (**Fig. 4C**), whereas cells transfected with the base editor alone retained a cytoplasmic distribution (0.25), confirming that rescue was sgRNA-dependent (**Fig. 5B**). Targeted amplicon sequencing of sorted cells confirmed A-to-G conversion at the p.C170Y position with an editing efficiency of 83.4% (**Fig. 5C**), which was independently verified by Sanger sequencing (**Fig. 5D**).

**Figure 5.**
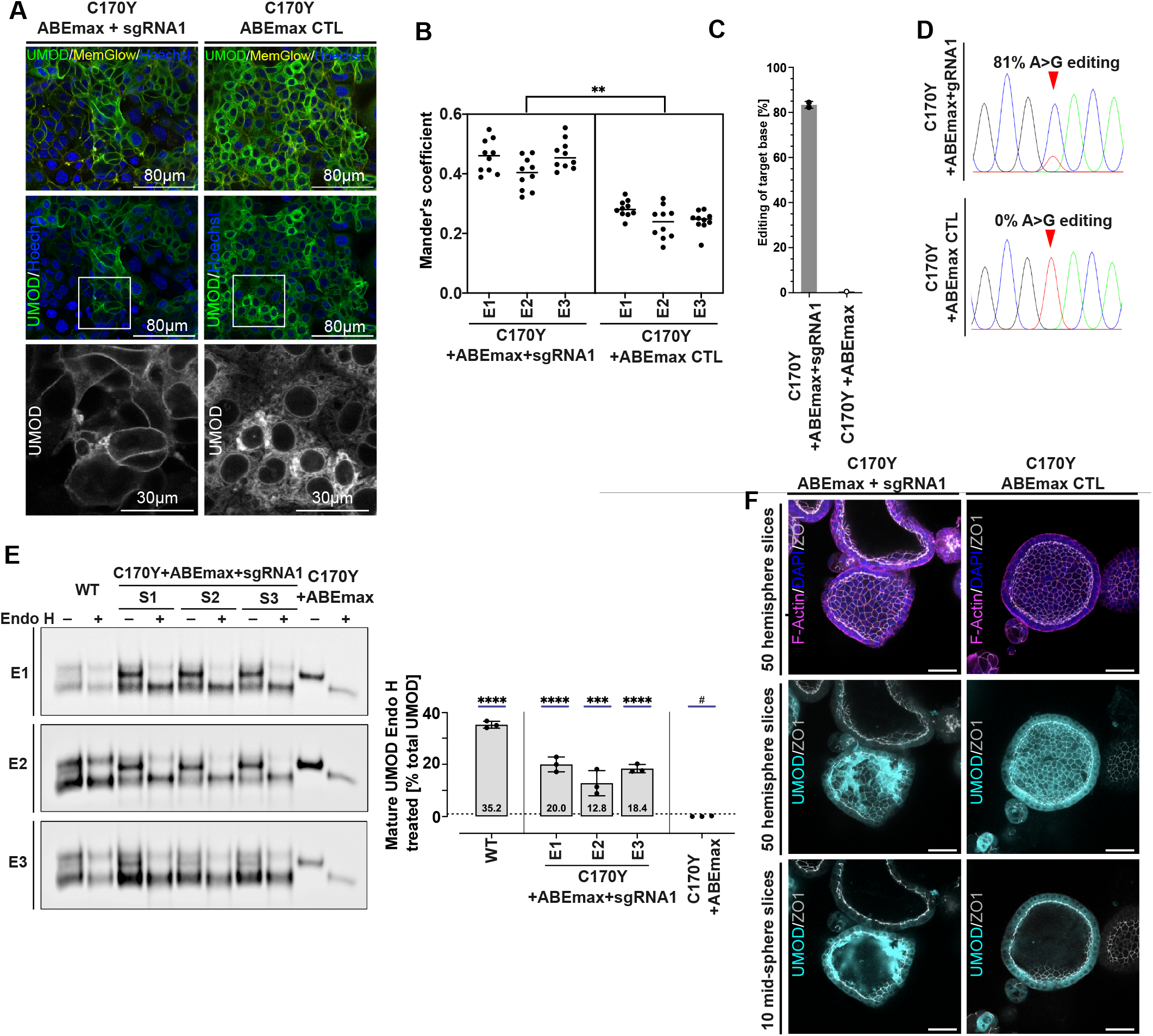
Base editing of the p.C170Y pathogenic variant restores uromodulin trafficking and maturation. (**A**) Representative confocal images of p.C170Y mutant hUMOD mIMCD-3 cells co-transfected with ABEmax-BFP and the sgRNA or with ABEmax-BFP alone, imaged after FACS enrichment of BFP-positive cells. GFP-tagged hUMOD (green), plasma membranes labeled with MemGlow (yellow), nuclei counterstained with Hoechst (blue). Scale bars, 80 µm (overview images) and 30 µm (magnified images). (**B**) Quantification of GFP-membrane colocalization by Manders’ coefficient for base-edited p.C170Y cells (C170Y + ABEmax+sgRNA1) and base editor only transfected p.C170Y cells (C170Y + ABEmax CTL). Data are shown as mean from n = 10 images (horizontal bars indicate mean values) across 3 independent biological experiments; dots represent data from individual images. Statistical significance was assessed using a two-tailed unpaired Student’s t-test. *\*\*P* < 0.01. (**C**) Editing efficiency at the p.C170Y target adenine in sorted cells, determined by TAS 72h after transfection. Data are shown as mean ± SD. (**D**) Representative Sanger sequencing traces showing A-to-G conversion at the p.C170Y target position in base-edited cells (C170Y + ABEmax+sgRNA1) and base editor only transfected cells (C170Y + ABEmax CTL). (**E**) Top: western blot analysis of UMOD in WT, C170Y +ABEmax+sgRNA1, and C170Y + ABEmax CTL mIMCD-3 cell lysates following mock treatment or Endo H digestion. Bottom: quantification of the Endo H-resistant, mature uromodulin fraction. Data are shown as mean ± SD from n = 3 replicates; ordinary one-way ANOVA with multiple comparisons, *\*\*\*P* < 0.001, *\*\*\*\*P* < 0.0001, # denotes the reference group for statistical comparisons. (**F**) Representative immunofluorescence images of 3D tubuloids expressing GFP-tagged p.C170Y hUMOD or p.C170Y hUMOD following electroporation with ABEmax mRNA and sgRNA1. Tubuloids were stained for zonula occludens-1 (ZO-1, light grey), F-actin (magenta), and nuclei (DAPI, blue), UMOD-GFP signal is shown in cyan. Panels show images from either 50 slices from a hemisphere or 10 slices mid-sphere, as indicated. Scale bars, 50 µm.

To confirm rescue at the protein level, lysates from WT, unedited and base-edited p.C170Y mIMCD-3 cells were digested with Endo H or PNGase F (**Fig. 5E**). In WT cells, a fraction of uromodulin was Endo H-resistant (35.2±1.3%), consistent with mature, Golgi-processed protein, whereas PNGase F removed all glycans. In unedited p.C170Y cells, uromodulin was fully Endo H-sensitive, indicating that all protein was retained in the ER in its immature form. Following base editing with ABEmax and C170Y_sgRNA1, a substantial fraction of mature, Endo H-resistant uromodulin of up to 20.0±2.8% was restored (n = 3) (**Fig. 5E**). Base editing of the p.C170Y variant therefore restores both uromodulin trafficking to the plasma membrane and maturation along the secretory pathway.

### Correcting p.C170Y in human 3D tubuloids

We next applied the leading combination identified in mIMCD-3 cells to the polarized human system. 3D tubuloids expressing p.C170Y hUMOD were electroporated with ABEmax mRNA together with C170Y_sgRNA1, or with ABEmax alone. Base-edited tubuloids showed a pronounced redistribution of uromodulin from intracellular compartments to the apical membrane, together with large luminal deposits resembling those observed in WT tubuloids and consistent with apical release of the corrected protein (**Fig. 5F**). No rescue was observed in tubuloids receiving the base editor alone. Correction of the p.C170Y variant therefore restores not only membrane trafficking but also the apical fate of uromodulin in a polarized human epithelium.

### Activating endogenous UMOD expression in primary human urine-derived cells

The lentiviral models described above express uromodulin from an intronless cDNA and cannot report strategies that act on the endogenous gene, such as splice-site disruption or nonsense-mediated decay of transcripts carrying a premature stop codon. We therefore established a CRISPR activation (CRISPRa) system to induce transcription from the endogenous *UMOD* locus. We employed the synergistic activation mediator (SAM) system^54^, in which a catalytically inactive Cas9 (dCas9) fused to the VP64 transcriptional activation domain acts together with the MS2-P65-HSF1 (MPH) activator helper complex, first implemented in a HEK293T line stably expressing the SAM components (**Fig. 6A**). To identify effective guides, we screened ten sgRNAs targeting the proximal *UMOD* promoter within 239 bp upstream of the transcription start site (act_sgRNA1–10; **Fig. 6B**), binding to both the sense- and the antisense strand and quantified *UMOD* mRNA by qRT-PCR (**Fig. 6C**). Individual guides increased *UMOD* expression between 10- and 1,000-fold relative to non-activated controls, and combining the three most effective guides (act_sgRNA1, act_sgRNA2 and act_sgRNA9, hereafter referred to as act_sgRNA mix) raised activation further, to up to 10,000-fold (**Fig. 6C**).

**Figure 6.**
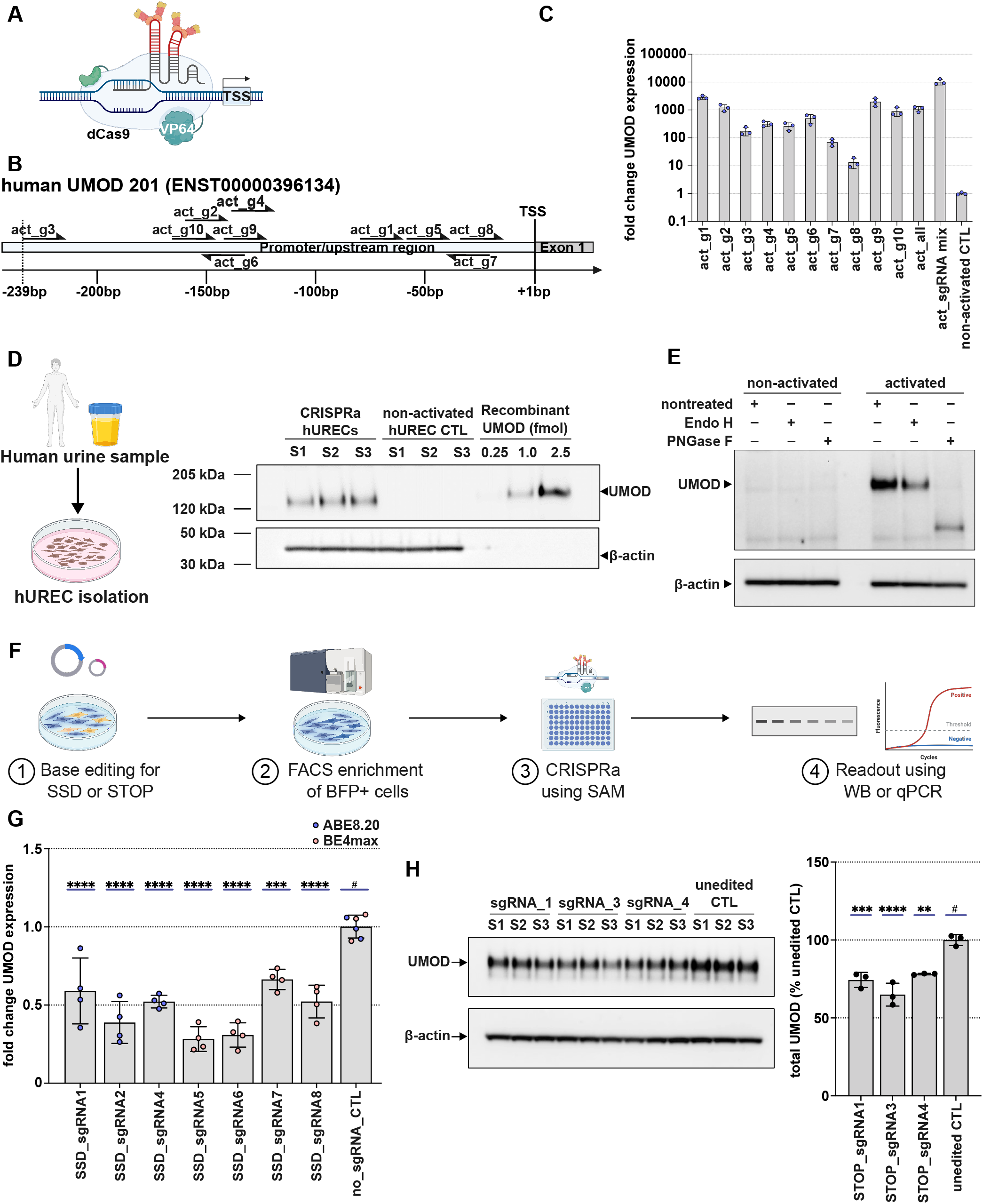
CRISPRa-mediated activation of endogenous UMOD enables functional testing of knockdown strategies. (**A**) Schematic of the synergistic activation mediator (SAM) system for CRISPR-mediated transcriptional activation. MS2-containing sgRNAs target the proximal UMOD promoter and recruit dCas9-VP64 together with the MS2-p65-HSF1 (MPH) helper complex. (**B**) Positions of the ten sgRNAs (act_sgRNA1–10) targeting the proximal UMOD promoter. Guides target both strands (arrows) within −20 to −239 bp upstream of the transcription start site (TSS), located within exon 1. (**C**) UMOD expression quantified by ΔΔCt qRT-PCR in CRISPRa-competent HEK293T cells (HEK293T/ dCas9-VP64 + MPH) after transfection with each individual sgRNA, with a combination of all sgRNAs (act_all) or with a combination of the three best-performing guides (act_sgRNA mix). CTL, mock transfection with non-targeting sgRNAs. Data are shown as mean ± SD from n = 3 replicates; experiment was independently repeated (n=2). (**D**) Left: schematic of hUREC isolation from human urine. Right: western blot of hUREC lysates after transfection with act_sgRNA mix or non-targeting sgRNAs, from three activated samples. Recombinant Uromodulin (0.25, 1 and 2.5 fmol) was loaded on the same blot for quantitative reference. β-actin served as loading control. (**E**) Deglycosylation analysis of CRISPRa-activated hURECs. Lysates were mock-treated or digested with Endo H or PNGase F. β-actin served as loading control. (**F**) Schematic of the experimental workflow: 1. base editing for splice-site disruption or stop codon introduction, 2. FACS enrichment of edited cells, 3. CRISPRa activation, and 4. analysis by qRT-PCR or western blot. **(G)** UMOD mRNA levels quantified by qRT-PCR in CRISPRa-activated HEK293T cells following splice-site disruption with seven sgRNA–base editor combinations (SSD_sgRNA1, 2, 4, 5, 6, 7 and 8 with ABE8.20m or BE4max), compared with base editor-only (no_sgRNA_CTL) controls (n = 4 biological replicates). (**H**) Left: Western blot of uromodulin in CRISPRa-activated HEK293T cells following stop codon introduction with STOP_sgRNA1, 3 and 4 and TadCBEd-V106W, compared with base editor-only transfected controls. Right: quantification of uromodulin protein levels (n = 3 biological replicates). β-actin served as loading control. (**G, H**) Data are shown as mean ± SD. Statistical significance was determined using ordinary one-way ANOVA followed by multiple comparisons testing. * *P* < 0.05, ** *P* < 0.01, *** *P* < 0.001, **** *P* < 0.0001; ns, not significant.

Having established robust activation in HEK293T cells, we asked whether endogenous *UMOD* could also be induced in primary human renal tubular epithelial cells. We isolated hURECs from spot urine samples of healthy donors and applied the CRISPRa system in 2D culture, detecting endogenous uromodulin protein by western blot in three independently activated hUREC cultures, whereas no protein was detectable in non-activated cells (**Fig. 6D**). Comparison with a recombinant uromodulin standard run on the same blot indicated that activated cultures produced uromodulin in the low femtomole range (**Fig. 6D**). To determine whether the induced protein was correctly processed, we again applied Endo H and PNGase F digestion (**Fig. 6E**). Activated hURECs yielded Endo H-resistant uromodulin that was fully deglycosylated by PNGase F, indicating passage through the Golgi and acquisition of mature complex glycans. CRISPRa therefore restores expression of correctly processed endogenous uromodulin in primary human renal epithelial cells obtained non-invasively from urine, providing a system in which base editing strategies acting on the endogenous *UMOD* transcript can be functionally assessed.

### Base editing reduces endogenous UMOD expression in CRISPRa-activated cells

We next used this system to test whether the inactivation strategies established at the DNA level reduce endogenous *UMOD* expression. HEK293T/dCas9-VP64+MPH cells were transfected with base editors and sgRNAs, enriched for successfully transfected cells by FACS, and subsequently activated with act_sgRNA_mix before analysis by RT-qPCR or western blot (**Fig. 6F**). Seven sgRNA-base editor combinations targeting essential splice sites (SSD_sgRNA1, 2, 4, 5, 6, 7 and 8 with ABE8.20m or BE4max) all reduced *UMOD* mRNA levels relative to activated cells transfected with base editor alone, demonstrating effective knockdown of the endogenous transcript through splice-site disruption. (**Fig. 6G**).

To assess stop codon introduction at the protein level, we focused on the three base editor STOP_sgRNA combinations closest to the translation start codon achieving editing efficiencies of above 50% (STOP_sgRNA1, STOP_sgRNA3 and STOP_sgRNA4). We analyzed uromodulin by western blot following editing with TadCBEd-V106W, which resulted in reduced uromodulin protein relative to activated cells transfected with base editor alone (**Fig. 6H**). CRISPRa-mediated activation of endogenous *UMOD* therefore provides a platform for functional evaluation of loss-of-function base editing strategies, validated here for both splice-site disruption and premature stop codon introduction.

## Discussion

ADTKD-*UMOD* is a toxic gain-of-function ER storage disease for which no approved disease-modifying treatment exists. Here we systematically evaluate base editing at the UMOD locus, from direct correction of a pathogenic missense variant to four mutation-agnostic strategies for lowering uromodulin. Correction of the p.C170Y variant restored uromodulin trafficking and maturation in a murine tubular cell line and in polarized human 3D tubuloids, establishing that the cellular disease phenotype is reversible by a single base conversion. To assess strategies acting on the endogenous gene, we developed a CRISPR activation platform that induces transcription from the endogenous UMOD locus in primary human cells.

As ADTKD-*UMOD* is driven by a toxic gain-of-function rather than by loss of protein, it is not amenable to gene replacement. Conventional Cas9-mediated correction depends on homology-directed repair, which is inefficient in the largely quiescent tubular epithelium and carries the risk of unintended insertions or deletions. Base editing avoids both limitations by installing single-nucleotide changes without double-strand breaks or donor templates^48,49^, making it well suited to the predominantly missense *UMOD* variant spectrum. We achieved efficient on-target correction of the p.C170Y variant, exemplifying the cysteine-disrupting missense mutations frequently located in exon 3, with minimal bystander editing. Correction restored both uromodulin trafficking to the plasma membrane and acquisition of mature glycans. Independent evidence indicates that the endpoints rescued here predict therapeutic benefit. In previous studies, relieving ER retention pharmacologically improved disease features in *Umod* knock-in mice. MANF-stimulated clearance of mutant uromodulin reduced fibrosis and inflammatory infiltration and improved TAL function^55^. More recently, TMED cargo receptors were shown to trap mutant uromodulin in early secretory compartments, and their small-molecule targeting restored apical localization and reduced ER stress and fibrosis^56^. Base-editing mediated correction acts further upstream than these approaches. It prevents synthesis of the misfolded protein and may restore physiological levels of wild-type uromodulin, rather than clearing or rerouting the mutant protein after translation. In principle, a single intervention would replace lifelong treatment.

As not all *UMOD* variants are directly correctable by base editing, mutation-agnostic strategies extend the approach to a broader range of patients. Several lines of evidence support *UMOD* inactivation as a therapeutic principle. *Umod*-deficient mice show no ADTKD phenotype^57^, although they are more susceptible to urinary tract infections^7^. Population sequencing likewise provides no evidence that *UMOD* is intolerant to loss-of-function variation^19^. Most directly, allele-selective antisense oligonucleotides were recently shown to lower immature uromodulin and reduce tubulointerstitial damage in *Umod* knock-in mice^58^, indicating that lowering mutant uromodulin can modify the disease in vivo.

We show that premature stop codon introduction and splice-site disruption can both be achieved efficiently at the *UMOD* locus. Building on these findings, further characterization of transcript processing and protein products will help establish the therapeutic potential of these approaches and identify potential escape mechanisms. By now, splice-site disruption has reached clinical testing, where adenine base editing of the PCSK9 splice donor site lowers LDL cholesterol after a single *in vivo* dose^59^, while stop codon introduction has been applied in human CAR T cells *ex vivo*^60^.

Promoter editing provides a third mutation-agnostic option, with a distinct safety rationale. Converting the common rs4293393-A risk allele to the protective G allele introduces no novel sequence but recreates a haplotype carried by approximately 19% of Europeans and associated with reduced *UMOD* transcription and protection against CKD^23,61^. Notably, the G allele is also underrepresented on pathogenic *UMOD* haplotypes, suggesting a potential disease-modifying effect in ADTKD-*UMOD* that remains to be established^61^.

A similar approach has been explored for sickle cell disease through base editing of the *HBG1* and *HBG2* promoters to recreate variants associated with hereditary persistence of fetal hemoglobin^62^. Promoter editing may enable partial attenuation of *UMOD* expression, potentially offering an advantage if complete inactivation proves undesirable.

One important consideration for inactivation strategies is the potential protective role of the WT allele. Deletion of the WT *Umod* allele in a heterozygous *Umod*^*R185S*^ knock-in mouse model increased uromodulin aggregate formation and tubular damage, though without affecting clinical parameters^35^. This raises the question of whether non-allele-specific inactivation could be detrimental in the long term. We therefore explored allele-specific stop codon introduction, exploiting the sequence alteration introduced by the p.R185S variant to restrict editing to the mutant allele. Promoter editing may offer a further route to allele selectivity. In patients heterozygous at rs4293393-A in cis with the pathogenic variant, only the mutant allele would be targeted. This selective approach preserves WT uromodulin production and may be preferable for clinical translation, though its generalizability across the broader *UMOD* variant spectrum will require case-by-case assessment.

This study establishes two human model systems that address complementary limitations, one providing access to the endogenous UMOD locus and one providing a human polarized epithelium. CRISPRa induces transcription from the endogenous locus, allowing loss-of-function strategies to be tested in cells that do not otherwise express uromodulin. Unlike cDNA overexpression, it preserves the introns and exon-exon junctions required to assess splice-site disruption and nonsense-mediated decay. The platform is equally applicable to antisense oligonucleotides or small molecules acting on the endogenous transcript, and transferable to other loci lacking a physiologically expressing cell model. The 3D tubuloid system addresses the second limitation of a functionally relevant read-out as the amount of urinary UMOD levels have been shown to negatively correlate with the ADTKD-phenotype. Uromodulin is apically sorted and released into the tubular lumen, a fate that cannot be observed in adherent culture, making tubuloids suited to interventions acting on protein folding and trafficking. Uromodulin is expressed here from a transgene in a mixed-segment epithelium rather than from the endogenous locus in the thick ascending limb and combining tubuloid culture with endogenous activation remains an objective for future work.

No editing was detected at the highest-ranked off-target sites predicted for the p.C170Y locus in silico, but genome-wide specificity remains a central consideration when selecting a lead combination. PAM-relaxed Cas9 variants engage more genomic sites than NGG-dependent SpCas9, and highly active deaminases may additionally mediate Cas9-independent deamination of DNA and RNA. Optimized base editors mitigate this trade-off, as TadCBEd-V106W, engineered for reduced off-target activity, matched or exceeded BE4max at most stop codon target sites. Unbiased genome-wide profiling, including assessment of Cas9-independent deamination, will be required before clinical translation. This study does not address *in vivo* delivery, vector immunogenicity or long-term genomic safety, all of which will require investigation in *Umod* knock-in models. Renal delivery remains the principal barrier, although capsid engineering is steadily improving tubular transduction, with context-dependent capsid selection and cross-species capsid evolution both yielding variants with enhanced renal tropism^63–65^, and local delivery routes offer a further option for nephron-restricted targeting.

In summary, base editing provides both variant-specific correction and mutation-agnostic silencing at the UMOD locus, and the CRISPRa platform established here extends beyond base editing to any intervention targeting endogenous UMOD. Establishing efficient delivery to the thick ascending limb and defining the editing threshold required for therapeutic benefit are the principal remaining steps towards translation.

## Data Availability

The data supporting the findings of this study are available within the article and its Supplementary Data. Raw targeted amplicon sequencing data and microscopy images, together with associated imaging metadata, have been deposited in the NCBI Sequence Read Archive (SRA) and Zenodo and will be made available upon publication. Additional source data underlying the figures, including quantitative PCR data and processed sequencing data, are provided in the Supplementary Data and accompanying Source Data files. Further information is available from the corresponding authors upon request.

## Acknowledgements

The figures were created with BioRender.com. During the preparation of this work the authors used AI-assisted tools (OpenAI GPT-5 mini and Claude Opus 5) to improve readability and language. After using this tool, the authors reviewed and edited the content as needed and take full responsibility for the content of the publication.

## Author contributions Statements

Conceptualization V.K., M.M.K, Data curation N.D., S.W., A. AS., G.Y., R.S., E.H., S.K., C.C., D.P., A.F., B.R., Formal analysis N.D., S.W., A. AS., G.Y., R.S., E.H., S.K., C.C., Funding acquisition V.K., M.M., Investigation V.K., M.M.K, Methodology V.K., M.M.K, O.D., J.H., Project administration V.K., M.M.K, Resources V.K., M.M.K, K.U.E, O.D., A.F., B.R., J.H., M.M., R.D., Supervision V.K., M.M.K, Validation N.D., Visualization N.D., V.K., Writing original draft N.D., V.K., M.M.K., Writing-review & editing all authors

## Funding

This work was supported by the Deutsche Forschungsgemeinschaft DFG (KL 3224/4-1 to V.K.; KA5060/5-1 to M.M.K.; HA6909/4-1 to J.H.; 539950728 to R.S.). V.K. was supported by the Else Kröner-Fresenius-Stiftung (2024_EKEA.34). M.M.K. was supported by the Emmy-Noether Program of the DFG (KA 5060/1-1). V.K. and M.M.K. were supported by the BIH Charité Clinician Scientist Program funded by the Charité – Universitätsmedizin Berlin and the Berlin Institute of Health at Charité. N.D. was supported by the Peter-Scriba MD stipend of the DGIM. K.U.E. and J.H. are supported by the European Program for Rare Diseases (EJP RD) for the establishment of an ADTKD-Net, a European patient registry (01GM2402A). A. F. was funded by the Urological Research Foundation Berlin. O.D. and M.M. are supported by the ADTKD-Net consortium of the European Joint Program on Rare Diseases; the Swiss National Science Foundation (grant 10.003.608); the University Research Priority Program (URPP) ITINERARE at the University of Zurich; and the Fondation Saint-Luc at UCLouvain (Brussels, Belgium).

## Conflicts of interest

A patent application related to this work has been filed. M.M.K. serves as a consultant for Insmed. M.M.K. is a co-inventor on patent applications related to gene editing and diagnostic technologies filed by Charité – Universitätsmedizin Berlin, the Max Delbrück Center for Molecular Medicine, and the Massachusetts Institute of Technology.

