## Supplementary Materials for "Base Editing as a Therapeutic Strategy for Autosomal Dominant Tubulointerstitial Kidney Disease caused by Pathogenic *UMOD* Variants"

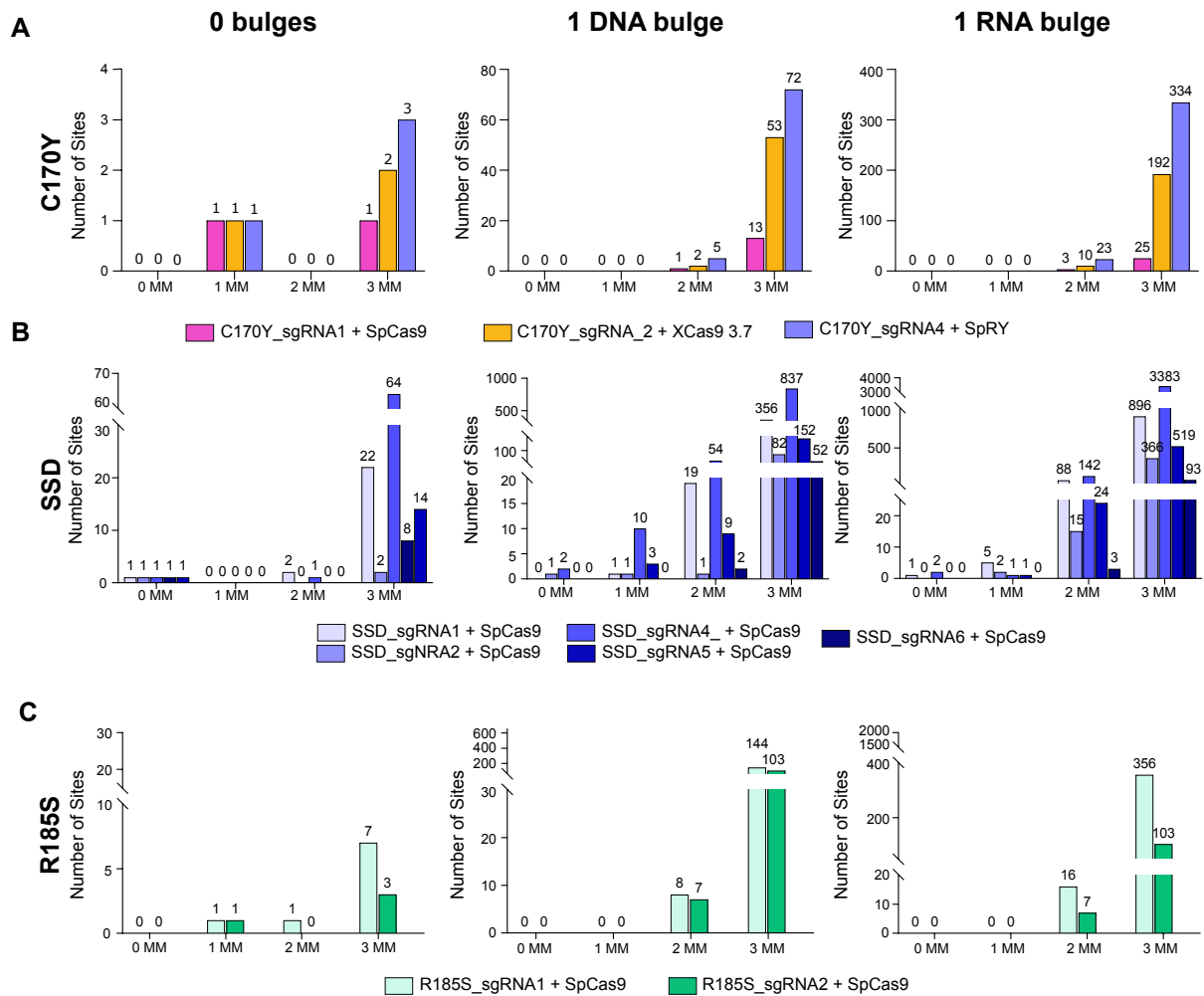

**Supplementary Figure S1.** In silico prediction of off-target sites for UMOD C170Y variant correction, splice-site disruption (SSD), and R185S-specific stop codon introduction. **(A–C)** sgRNA- and PAM-dependent off-target sites were identified computationally using the web-based tool Cas-OFFinder. Sites with up to three mismatches (MM) and either no bulge, a 1-nt DNA bulge, or a 1-nt RNA bulge relative to the sgRNA sequence were counted as potential off-target sites. Y-axes show the numbers of predicted off-target sites, while x-axes indicate the numbers of mismatches. Bars are color-coded according to the respective sgRNA and Cas9 variant combinations, as indicated in the legends. The numbers of predicted off-target sites are indicated above the bars. **(A)** For correction of the C170Y variant, C170Y\_sgRNA1 was evaluated in combination with SpCas9 (NGG PAM), C170Y\_sgRNA2 with xCas9 3.7 (NG PAM), and C170Y\_sgRNA4 with SpRY (NNN PAM). **(B)** For splice-site disruption (SSD), SSD\_sgRNAs 1, 2, 4, 5, and 6 were evaluated in combination with SpCas9 (NGG PAM). **(C)** For allele-specific stop codon introduction targeting the R185S variant, R185S\_sgRNA1 and R185S\_sgRNA2 were evaluated in combination with SpCas9 (NGG PAM).

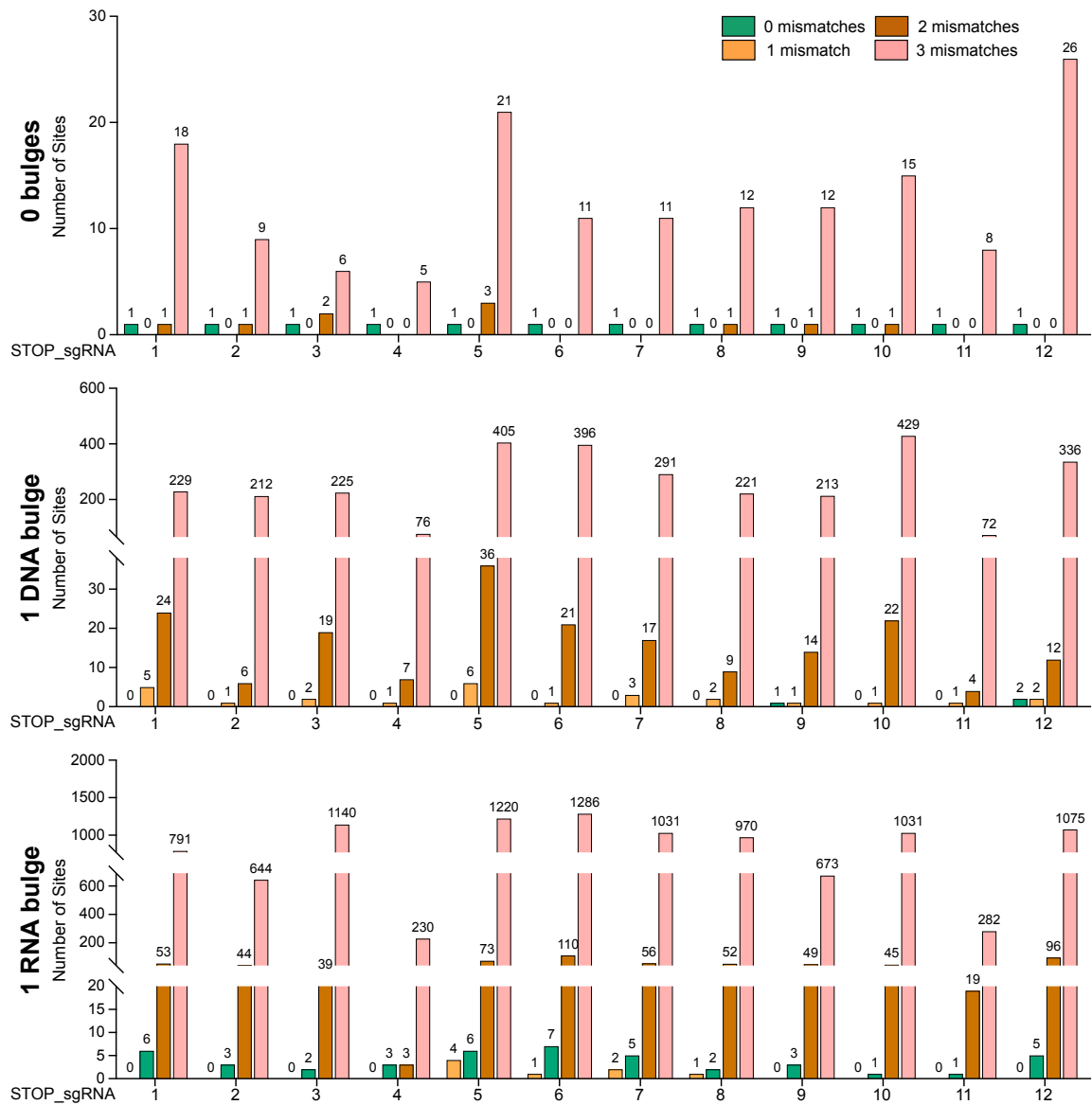

**Supplementary Figure S2.** In silico prediction of off-target sites for UMOD stop codon introduction. sgRNA- and PAM-dependent off-target sites were identified computationally using the web-based tool Cas-OFFinder. Sites with up to three mismatches and either no bulge, a 1-nt DNA bulge, or a 1-nt RNA bulge relative to the sgRNA sequence were counted as potential off-target sites. Y-axes show the number of predicted off-target sites, while the x-axes indicate the STOP\_sgRNAs screened. Bars are color-coded according to the number of mismatches, as indicated in the legend. The numbers of predicted off-target sites are indicated above the bars. For stop codon introduction, all sgRNAs were evaluated in combination with SpCas9 (NGG PAM).

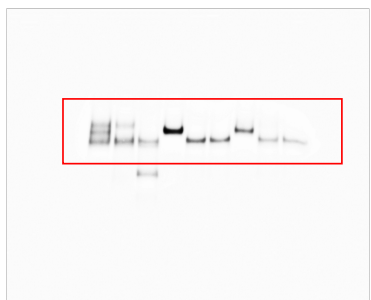

**Supplementary Figure S3.** Uncropped Western blots of UMOD deglycosylation analysis using Endo H and PNGase F treatment in mIMCD-3 cells according to **Figure 4D**. Box indicates the region shown in the main figure.

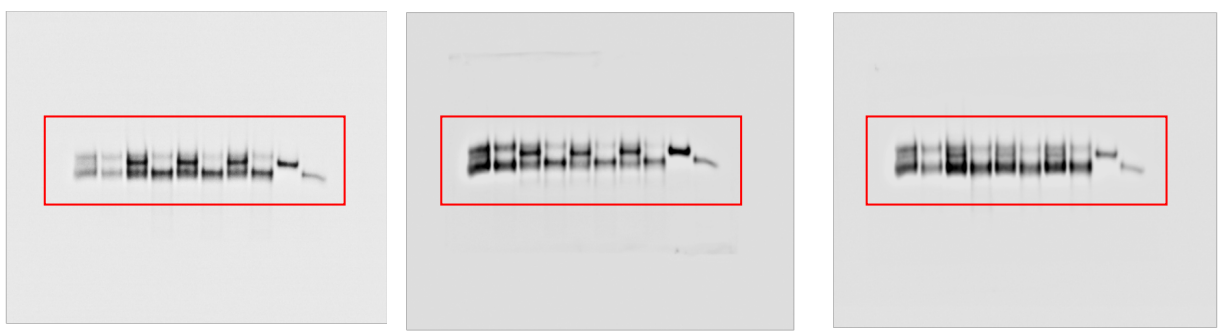

**Supplementary Figure S4.** Uncropped Western blots of UMOD deglycosylation analysis using Endo H treatment in WT, unedited and base-edited p.C170Y mIMCD3 cells as shown in **Figure 5E** (from left to right shown as top, middle and bottom). Boxes indicate the region shown in the main figure.

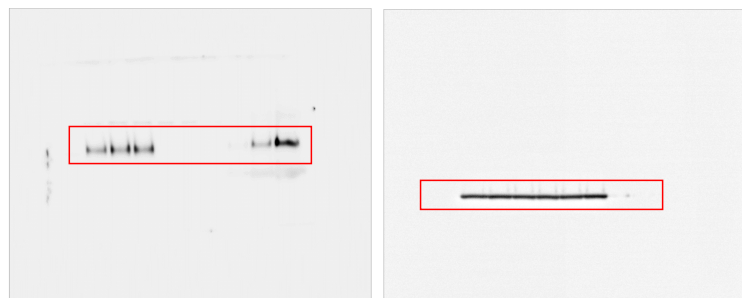

**Supplementary Figure S5.** Uncropped Western blots corresponding to **Figure 6D**. Left: Uncropped Western blot of UMOD in huREC lysates following transfection with the act\_sgRNA mix or non-targeting sgRNAs. Right: Uncropped Western blot of β-actin used as a loading control. Boxes indicate the regions shown in the main figure.

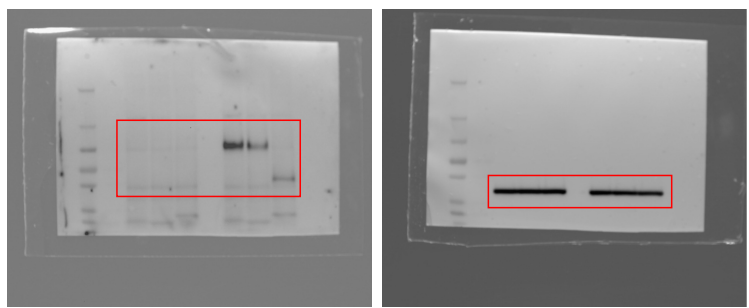

**Supplementary Figure S6.** Uncropped Western blots corresponding to **Figure 6E**. Left: Uncropped Western blot of UMOD in CRISPRa-activated huREC lysates following deglycosylation with Endo H or PNGase F. Right: Uncropped Western blot of β-actin used as a loading control. Boxes indicate the regions shown in the main figure.

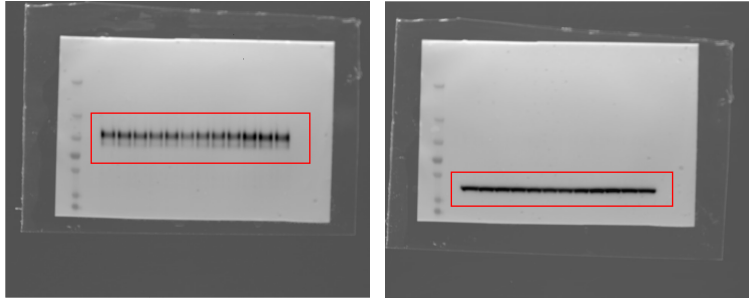

**Supplementary Figure S7.** Uncropped Western blots corresponding to **Figure 6H**. Left: Uncropped Western blot of UMOD in CRISPRa-activated HEK293T cells following stop codon introduction with STOP\_sgRNA1, 3 and 4 and TadCBEd-V106W, compared with base editor-only transfected controls. Right: Uncropped Western blot of β-actin used as a loading control. Boxes indicate the regions shown in the main figure.

**Supplementary Table S1: sgRNA sequences**

| **sgRNAs C170Y HEK293T screening** | **Sequence** |
| --- | --- |
| C170Y_sgRNA1 | CGTGTaCGCGGATCCGTGCC |
| C170Y_sgRNA2 | GTGTaCGCGGATCCGTGCCA |
| C170Y_sgRNA3 | TGTaCGCGGATCCGTGCCAG |
| C170Y_sgRNA4 | TCGTGTaCGCGGATCCGTGC |
| **sgRNAs hUMOD STOPs** | **Sequence** |
| STOP_sgRNA1 | TGGGGcAGCCATCTCTGACT |
| STOP_sgRNA2 | ACCTGTcAGGAGGGCTTCAC |
| STOP_sgRNA3 | GTGCcAGGCGCACCGCACCC |
| STOP_sgRNA4 | GCTGCGcCAGTACTCGTCCA |
| STOP_sgRNA5 | GGGCcAGGGCGGTGCGCGCA |
| STOP_sgRNA6 | GTCcAGGTGAAGGCCTGTGC |
| STOP_sgRNA7 | AAAcAGGACTTCAACATCAC |
| STOP_sgRNA8 | AAGTGCcAGCTGAAGAGTCT |
| STOP_sgRNA9 | CTAcAGCCAATGGTCAGGTG |
| STOP_sgRNA10 | ATCcAGGACAGGTAAGGCAA |
| STOP_sgRNA11 | GTCcAGATGTTCCGGTTTGC |
| STOP_sgRNA12 | GTGTCcAGGCCACAGTCTCA |
| **sgRNAs hUMOD SSDs** | **Sequence** |
| SSD_sgRNA1 | AGCCTTAcCTGAAGCCAGAA |
| SSD_sgRNA2 | GCTGAcCTGTGCAGTACGCC |
| SSD_sgRNA3 | CGGCTGAcCTGTGCAGTACG |
| SSD_sgRNA4 | CCCTGCaGACCCCAGCTCCG |
| SSD_sgRNA5 | GATATcTGAAACAGGTTAGG |
| SSD_sgRNA6 | ATATcTGAAACAGGTTAGGT |
| SSD_sgRNA7 | TTCCTcTGTTGCAGGGAATg |
| SSD_sgRNA8 | TCCTcTGTTGCAGGGAATgg |
| SSD_sgRNA9 | ATTCCTcTGTTGCAGGGAAT |
| SSD_sgRNA10 | CCTGGCaGTGCTCTAAACAT |
| SSD_sgRNA11 | gCCTTaCCTGTCCTGGATGA |
| SSD_sgRNA12 | CCTGTaGACCTGCTCTGGGA |
| SSD_sgRNA13 | CTCTTAcCTTTCCGTGTGAT |
| SSD_sgRNA14 | TCTCTTAcCTTTCCGTGTGA |
| **sgRNAs mIMCD-C170Y correction** | **Sequence** |
| mIMCD_sgRNA1 | CGTGTaCGCGGATCCGTGTC |
| **sgRNAs rs4293393 editing** | **Sequence** |
| rs4293393_sgRNA1 | AGTTCAGACATCACTGGACC |
| rs4293393_sgRNA2 | GTTCAGACATCACTGGACCT |
| **sgRNAs R185S Allele Specific STOP** | **Sequence** |
| R185S_sgRNA1 | TGCTGCTCCAGTACTCGTCC |
| R185S_sgRNA2 | GCTGCTcCAGTACTCGTCCA |
| **sgRNAs CRISPRa UMOD** | **Sequence** |
| ACT_sgRNA1 | CAGAACTAGAGACTAATTGG |
| ACT_sgRNA2 | ACAACCTCACAGCTTGGACA |
| ACT_sgRNA3 | TGTATTTCCGGGCACAGGTG |
| ACT_sgRNA4 | AAGGCAAACATTATGCCAGG |
| ACT_sgRNA5 | GAGGAGAGATTGCCAGCCTG |
| ACT_sgRNA6 | TTTGCCTTGTCCAAGCTGTG |
| ACT_sgRNA7 | TATATATACACATTTGCCCC |
| ACT_sgRNA8 | AATGTGTATATATAAGTATG |
| ACT_sgRNA9 | GACAAGGCAAACATTATGCC |
| ACT_sgRNA10 | ACAACAACAACCTCACAGCT |

**Supplementary Table S2: Base editors used and corresponding Addgene references**

| Base editor | Cas variant | Construct name | Source | Reference |
| --- | --- | --- | --- | --- |
| BE4max | SpCas9 | pCMV_BE4max | Addgene #112093 | Koblan et al., 2018; gift from David Liu |
| TadCBE d-V106W | SpCas9 | SpCas9 TadCBE d-V106W | Addgene #193839 | Neugebauer et al., 2023; gift from David Liu |
| CBE6b | SpCas9 | SpCas9-CBE6b | Addgene #215820 | Zhang et al., 2024; gift from David Liu |
| ABEmax | SpCas9 | pCMV_ABEmax | Addgene #112095 | Koblan et al., 2018; gift from David Liu |
| ABE8.20m | SpCas9 | ABE8.20-m | Addgene #136300 | Gaudelli et al., 2020; gift from Nicole Gaudelli |
| ABE8e | SpCas9 | ABE8e | Addgene #138489 | Richter et al., 2020; gift from David Liu |
| ABEmax | SpCas9-NG | pCMV-T7-ABEmax(7.10)-SpCas9-NG-P2A-EGFP (RTW4564) | Addgene #140005 | Walton et al., 2020; gift from Benjamin Kleinstiver |
| ABEmax | SpG | pCMV-T7-ABEmax(7.10)-SpG-P2A-EGFP (RTW4562) | Addgene #140002 | Walton et al., 2020; gift from Benjamin Kleinstiver |
| ABEmax | SpRY | pCMV-T7-ABEmax(7.10)-SpRY-P2A-EGFP (RTW5025) | Addgene #140003 | Walton et al., 2020; gift from Benjamin Kleinstiver |
| ABE8e | SpCas9-NG | pCMV-T7-ABE8e-nSpCas9-NG-P2A-EGFP (KAC999) | Addgene #185914 | Gift from Benjamin Kleinstiver (unpublished) |
| ABE8e | SpG | pCMV-T7-ABE8e-nSpG-P2A-EGFP (KAC984) | Addgene #185911 | Alves et al., 2024; gift from Benjamin Kleinstiver |
| ABE8e | SpRY | pCMV-T7-ABE8e-nSpRY-P2A-EGFP (KAC1069) | Addgene #185912 | Alves et al., 2024; gift from Benjamin Kleinstiver |
| ABE8.20m | SpCas9-NG | pCMV-T7-ABE8.20m-nSpCas9-NG-P2A-EGFP (KAC1164) | Addgene #185919 | Gift from Benjamin Kleinstiver (unpublished) |
| ABE8.20m | SpG | pCMV-T7-ABE8.20m-nSpG-P2A-EGFP (KAC1160) | Addgene #185916 | Alves et al., 2024; gift from Benjamin Kleinstiver |
| ABE8.20m | SpRY | pCMV-T7-ABE8.20m-nSpRY-P2A-EGFP (KAC1335) | Addgene #185917 | Alves et al., 2024; gift from Benjamin Kleinstiver |
| ABE8e | SaKKH | SaKKH-ABE8e | Addgene #138502 | Richter et al., 2020; gift from David Liu |
| ABE9 | SpG | pCD111_pCMV-T7-ABE9-SpG-P2A-tagBFP | Derived from Addgene #194208 | Chen et al., 2023; Walton et al., 2020; parental plasmid gift from Dali Li |
| ABE9 | SpRY | ABE9_nSpRY_tagBFP | Derived from Addgene #194208 | Chen et al., 2023; Walton et al., 2020; parental plasmid gift from Dali Li |
| ABEmax | SpCas9 | pCMV_ABEmax_P2A_tagBFP | Derived from Addgene #112101 | Koblan et al., 2018; parental plasmid gift from David Liu |
| ABE8.20m | SpCas9 | ABE8.20m_P2A_tagBFP | Derived from Addgene #136300 | Gaudelli et al., 2020; parental plasmid gift from Nicole Gaudelli |
| BE4max | SpCas9 | pCMV_BE4max_BFP | Derived from Addgene #112093 | Koblan et al., 2018; parental plasmid gift from David Liu |
| TadCBE d-V106W | SpCas9 | SpCas9 TadCBE d-V106W_tagBFP | Derived from Addgene #193839 | Neugebauer et al., 2023; parental plasmid gift from David Liu |

**Supplementary Table S3: Gibson assembly primers used to introduce tagBFP into base editor constructs**

| Primer | Sequence |
| --- | --- |
| GA_BFP_P1 | ACCCTGGACCTATGAGCGAGCTGATTAAGGAGAAC |
| GA_BFP_P2 | CTTCTGACAACGATCGGAGGACCGAAG |
| GA_BFP_P3 | GTCCTCCGATCGTTGTCAGAAGTAAGTTGG |
| GA_BFP_P4 | AGCTCGCTCATAGGTCCAGGGTTCTCCTCCA |
| GA_BFP_P6 | AACCACCAGAGTGCCCCAGTTTGCTAGGG |
| GA_BFP_P7 | ACTGGGGCACTCTGGTGGTTCTCCCAAGAAGA |

**Supplementary Table S4: Gibson assembly primers used to clone synthesized target sequences into lentiviral or Sleeping Beauty vectors**

| Primer | Sequence |
| --- | --- |
| GA_lenti_P1 | GTCCTCCGATCGTTGTCAGAAGTAAGTTGG |
| GA_lenti_P2 | CTTCTGACAACGATCGGAGGACCGAAG |
| GA_lenti_P3 | GATCTCGATAGCAGGCCTAACACATGCAAGTC |
| GA_lenti_P4 | CTAGCTACTAGAAGCAGTGGTATCAACGCAGAG |
| GA_lenti_P5 | GATACCACTGCTTCTAGTAGCTAGCTAGCTAGTCG |
| GA_lenti_P6 | GTTAGGCCTGCTATCGAGATCTGAGTCCGG |
| GA_SB_P1 | CTTCTGACAACGATCGGAGGACCGAAG |
| GA_SB_P2 | GTCCTCCGATCGTTGTCAGAAGTAAGTTGG |
| GA_SB_P3 | CAGATATCCATCGCAAATGGGCGGTAGGC |
| GA_SB_P4 | CTAGTGAATTCCGCTCACTCAAAGGCGGTAATCA |
| GA_SB_P5 | CTTTGAGTGAGCGGAATTCACTAGTGTCGACGAG |
| GA_SB_P6 | GCCCATTTGCGATGGATATCTGCAGGATCCGCC |

**Supplementary Table S5: PCR conditions for targeted amplicon sequencing (TAS)**

| PCR-1 |  |  |
| --- | --- | --- |
| Cycles | Temperature | Time |
| 1x | 98°C | 2min |
| 35x | 98°C | 10s |
|  | 68°C | 12s/20s |
|  | 72°C | 12s/20s |
| 1x | 72°C | 2min |

| PCR-2 |  |  |
| --- | --- | --- |
| Cycles | Temperature | Time |
| 1x | 98°C | 2min |
| 8-12x | 98°C | 10s |
|  | 65°C | 30s |
|  | 72°C | 30s |
| 1x | 72°C | 2min |

**Supplementary Table S6: qPCR primer and probe sequences**

| Primer Name | Sequence |
| --- | --- |
| UMOD_qPCR_fwd | GTGTCACCTGGCGTACTG |
| UMOD_qpcR_rev | TCCAGGAGGGAGATAT |
| UMOD_probe | CCTGTTTGCACTGGC |
| ACTB_qPCR_fwd | CCTTGCACATGCCGGAG |
| ACTB_qpcR_rev | ACAGAGCCTCGCCTTTG |
| ACTB_probe | GTGAGCTGGCGG |

**Supplementary Table S7: qPCR cycling conditions**

| Step | Temperature | Time | Cycles |
| --- | --- | --- | --- |
| Reverse Transcription | 55°C* | 10min | 1 |
| Initial Denaturation | 95°C | 1min | 1 |
| Denaturation | 95°C | 10s | 45 |
| Extension | 60°C | 30s (+ plate read) |  |
